# An insect serotonin receptor mediates cellular immune responses and its inhibition by phenylethylamide derivatives from bacterial secondary metabolites

**DOI:** 10.1101/556159

**Authors:** Ariful Hasan, Hyun-Suk Yeom, Jaewook Ryu, Helge B. Bode, Yonggyun Kim

## Abstract

Serotonin (5-hydroxytryptamine: 5-HT) is a biogenic monoamine that mediates immune responses and modulates nerve signal in insects. *Se-5HTR*, a specific receptor of serotonin, has been identified in the beet armyworm, *Spodoptera exigua.* It is classified into subtype 7 among known 5HTRs. *Se-5HTR* was expressed in all developmental stages of *S. exigua.* It was expressed in all tested tissues of larval stage. Its expression was up-regulated in hemocytes and fat body in response to immune challenge. RNA interference (RNAi) of *Se-5HTR* exhibited significant immunosuppression by preventing cellular immune responses such as phagocytosis and nodulation. Treatment with an inhibitor (SB-269970) specific to 5HTR subtype 7 resulted in significant immunosuppression. Such immunosuppression was also induced by bacterial secondary metabolites derived from *Xenorhabdus* and *Photorhabdus*. To determine specific bacterial metabolites inhibiting Se-5HTR, this study screened 37 bacterial secondary metabolites with respect to cellular immune responses associated with Se-5HTR and selected 10 potent inhibitors. These 10 selected compounds competitively inhibited cellular immune responses against 5-HT and shared phenylethylamide (PEA) chemical skeleton. Subsequently, 46 PEA derivatives were screened and resulting potent chemicals were used to design a compound to be highly inhibitory against Se-5HTR. The designed compound was chemically synthesized. It showed high immunosuppressive activities along with specific and competitive inhibition activity for Se-5HTR. This study reports the first 5HT receptor from *S. exigua* and provides its specific inhibitor designed from bacterial metabolites and their derivatives.

**Author Summary:** Serotonin (5-hydroxytryptamine: 5-HT) plays a crucial role in mediating nerve and immune signals in insects. Interruption of 5-HT signal leads to malfunctioning of various insect physiological processes. Se-5HTR, a 5-HT receptor of beet armyworm, *Spodoptera exigua,* was identified and classified as subtype 7 (5-HT_7_) of 5-HT receptors. A specific inhibitor (SB-269970) for 5-HT_7_ highly inhibited immune responses such as phagocytosis and nodulation mediated by Se-5HTR. Two entomopathogenic bacteria, *Xenorhabdus* and *Photorhabdus,* could secrete potent inhibitors against immune responses mediated by 5-HTR. Bacterial secondary metabolites were screened against Se-5HTR-mediating immune responses. Most of resulting compounds shared phenylethylamide (PEA) chemical skeleton. Subsequent screening using PEA derivatives supported the importance of this chemical skeleton. Based on their relative inhibitory activities, a compound was designed and synthesized. This novel compound possessed high inhibitory activities against Se-5HTR-mediating immune responses and exhibited competitive inhibition with 5-HT.

## Introduction

Serotonin or 5-hydroxytryptamine (5-HT) is a biogenic monoamine found across most phyla of life [1]. This indolamine compound is biosynthesized from tryptophan by successive catalytic activities of tryptophan hydroxylase and aromatic-L-amino acid decarboxylase [2–4] primarily in nervous systems [5,6]. In human and other vertebrates, serotonin is a well-known neurotransmitter involved in mood, appetite, sleep, anxiety, cognition, and psychosis [7–9]. Outside the nervous system, serotonin plays important roles as growth factor and regulator of hemostasis and blood clotting [10,11]. In plants, serotonin is basically involved in stress signaling [12]. Serotonin plays crucial role in physiological and behavioral processes in insects and other invertebrates [13,14]. In *Drosophila melanogaster*, there is evidence that serotonin is required for courtship and mating [15], circadian rhythm [16,17], sleep [18], locomotion [13,19], aggression [20], insulin signaling and growth [21], and phagocytosis [22]. Serotonin is also involved in olfactory processing [23], feeding behavior [19], heart rate [24], and responses to light [25] in *D. melanogaster* larvae.

Serotonin modulates physiological processes by binding to specific receptors. Seven main subtypes of serotonin receptors have been classified in vertebrates [26]. Except 5-HT_3_ receptor, the other six classes (5-HT_1_, 5-HT_2_, 5-HT_4_, 5-HT_5_, 5-HT_6_, and 5-HT_7_ receptors) belong to G protein-coupled receptor family [27]. Among these receptors, 5-HT_1_ and 5-HT_5_ receptors can inhibit cAMP synthesis by preferentially coupling to a trimeric G protein G_i/o_ [28]. 5-HT_2_ receptor uses G_q/11_ to induce breakdown of inositol phosphates, resulting in an increase in cytosolic Ca^2+^ level [28]. Besides, 5-HT_4_, 5-HT_6_, and 5-HT_7_ receptors coupled to G_s_ can stimulate cAMP production [28]. However, 5-HT_3_ receptor is a ligand-gated cation channel that mediates neuronal depolarization [29].

Insects have at least three subtypes of 5-HT receptors. Five different 5-HT receptors as orthologous to mammalian 5-HT_1A_, 5-HT_1B_, 5-HT_2A_, 5-HT_2B_, and 5-HT_7_ have been identified in *D. melanogaster* [14]. In addition, partial sequences of two 5-HT_1_, two 5-HT_2_, and one 5-HT_7_ receptors have been identified in the field cricket *Gryllus bimaculatus* [30]. Two 5-HT_1_ receptors and two 5-HT_1_ splice variants have been described in *Tribolium castaneum* and *Papilio xuthus,* respectively [31,32]. As from *D. melanogaster* and *G. bimaculatus,* two 5-HT_2_ receptors have been described from *Apis mellifera* while only one 5-HT_2_ receptor has been reported in other insects such as *Periplaneta americana* and *Locusta migratoria* [33,34]. In a lepidopteran species, *Pieris rapae,* four different 5-HT receptors including a novel subtype 8 have been reported [35,36].

5-HT modulates various physiological processes via expression of different receptor types in various tissues. 5-HT receptors are expressed highly in brain and ventral nerve cord of insects [32]. 5-HT_1_ receptor in honey bee brain is involved in visual information processing [37]. Expression pattern of 5-HT_7_ receptor in honey bee nervous system suggests its possible roles in information processing, learning, and memory [38]. 5-HT receptors might play roles in neuroendocrine secretion and gut motility in cockroaches [29]. In salivary gland of several insects, 5-HT_7_ receptor has been reported to be involved in salivary secretion mediated by cAMP level elevation [33,39,40]. Moreover, 5-HT_7_ receptor mediates visceral muscle contraction in the gastrointestinal tract of several insects including *A. aegypti* and *T. castaneum* [41,42]. 5-HT not only has neurophysiological roles, but also mediates cellular immune responses in insects by enhancing phagocytosis and nodulation [43]. Two different 5-HT receptors (1B and 2B) are expressed in hemocytes of *P. rapae,* of which 5-HT receptor 1B mediates cellular immune response [22]. In another lepidopteran insect, *Spodoptera exigua,* 5-HT mediates increase of total circulatory hemocyte number by stimulating sessile hemocytes and mediating cellular immune responses such as phagocytosis and nodule formation [44,45]. However, 5-HT receptor in *S. exigua* has not been reported yet.

Two entomopathogenic bacteria, *Xenorhabdus* and *Photorhabdus*, can inhibit insect immune responses to protect themselves and their symbiotic nematodes [46]. To accomplish host immunosuppression, these bacteria can synthesize and secrete secondary metabolites to inhibit immune signals and effectors [47]. Among these bacterial metabolites, tryptamine and phenylethylamide derivatives have been identified with suggested function of interrupting 5-HT signaling [48]. The objective of the present study was to determine bacterial secondary compound(s) that could inhibit 5-HT signaling. To this end, we identified 5-HT receptor that could mediate insect immunity in *S. exigua*. To determine a potent inhibitor of this receptor, we screened bacterial secondary metabolites and their potent derivatives.

## Results

### Bioinformatics analyses reveal that *S. exigua* contains a 5-HT receptor

From a short read archive database (GenBank accession number: SRR1050532) of *S. exigua,* a highly matched contig (accession number: GARL01017386.1) containing 3,308 bp nucleotide sequence with an open reading frame (ORF) from 996^th^ to 2,690^th^ bp was identified. Prediction of amino acid sequence by BlastP analysis revealed that its ORF sequence shared identities with other insect 5-HT receptors: 99% with *Spodoptera litura* 5HTR (XP_022827337.1), 98% with *Helicoverpa armigera* 5HTR (XP_021195909.1), 95% with *Trichoplusia ni* 5HTR (XP_026729862.1), and 85% with *Manduca sexta* 5HTR (AGL46976.1). This novel 5-HT receptor from *S. exigua* (Se-5HTR) encoded a sequence of 564 amino acids having a predicted molecular weight of about 63.23 kDa. Phylogenetic analysis of its protein sequence indicated that Se-5HTR was clustered with other 5-HT_7_ receptors (Fig 1A). Predicted amino acid sequence of Se-5HTR contained a signal peptide of 35 residues and attained GPCR character with seven transmembrane domains. Consensus N-linked glycosylation sites are located in the N-terminus (Asn^48^ and Asn^53^), the third intracellular loop (Asn^362^), and C-terminus (Asn^534^). Several consensus sites for phosphorylation by protein kinase A and/or protein kinase C were predicted throughout the length of the sequence. A disulfide bond between two Cys residues at extracellular matrix (‘ECM’, S1 Fig) is also predicted.

**Fig. 1.**
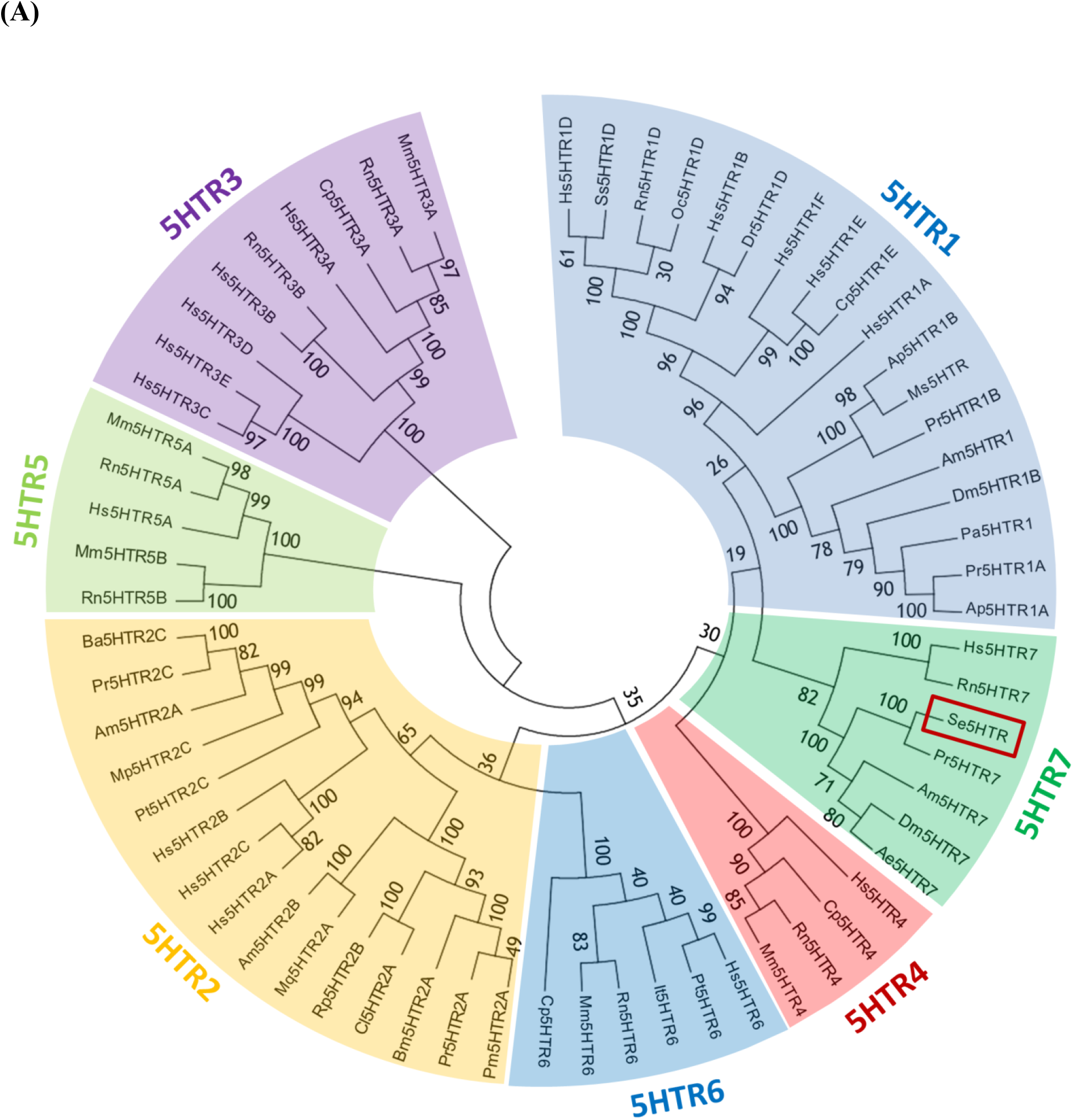
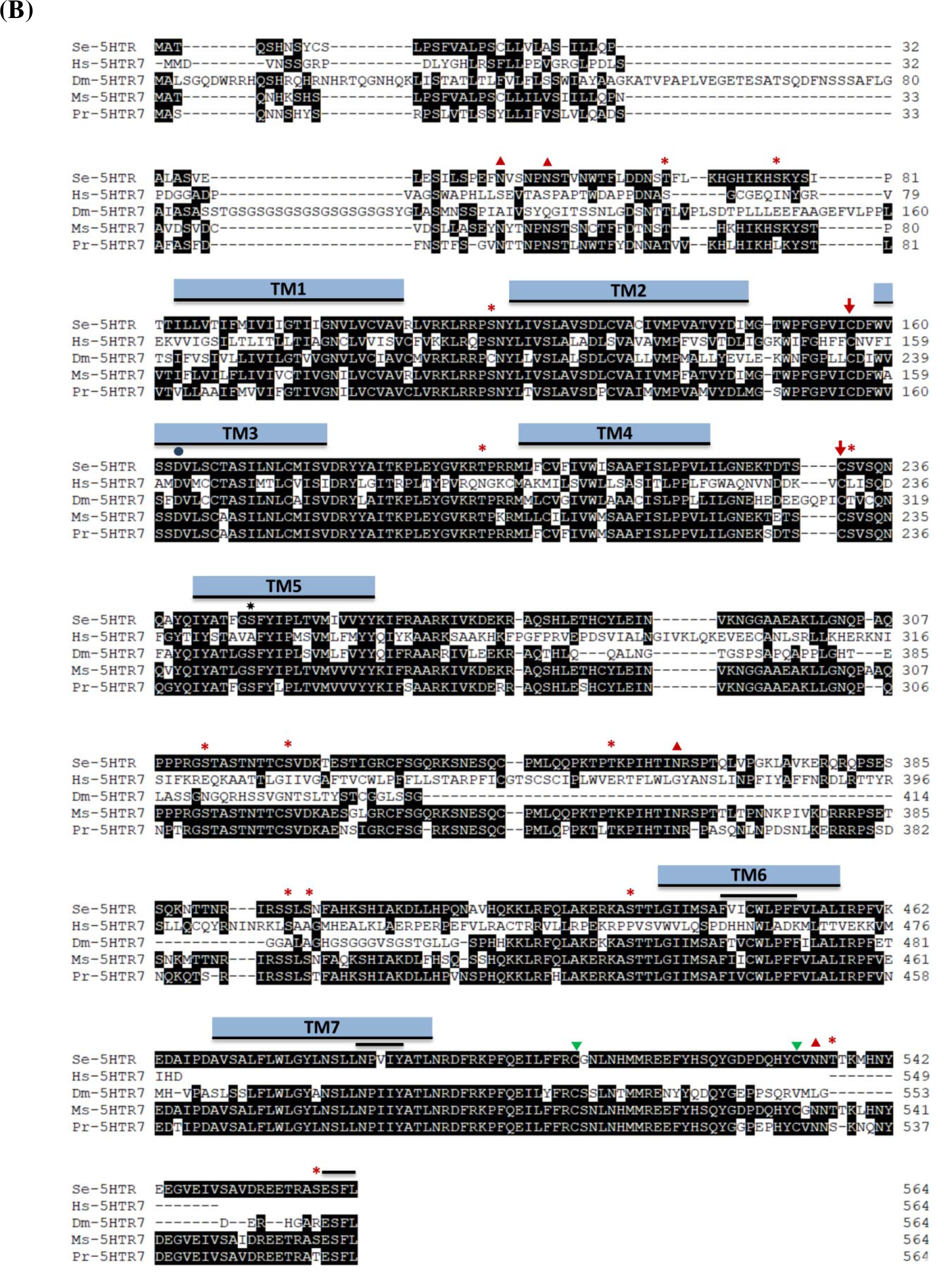
Bioinformatics analysis of a novel 5-HTR_7_ identified from *S. exigua* (Se-5HTR: MH025798). **(A)** Phylogenetic analysis of 5HTRs. Amino acid sequences were retrieved from GenBank: Hs5HTR1D (NP_000855.1), Ss5HTR1D (NP_999323.1), Rn5HTR1D (NP_036984.1), Oc5HTR1D (NP_001164624.1), NP_000854.1 Hs5HTR1B (NP_000854.1), Dr5HTR1D (NP_001139158.1), Hs5HTR1F (NP_000857.1), Hs5HTR1E (NP_000856.1), Cp5HTR1E (NP_001166222.1), Hs5HTR1A (NP_000515.2), Ap5HTR1B (ABY85411.1), Ms5HTR (ABI33827.1), Pr5HTR1B (XP_022120028.1), Am5HTR1 (NP_001164579.1), Dm5HTR1B (NP_001163201.2), Pa5HTR1 (CAX65666.1), Pr5HTR1A (XP_022129638.1), Ap5HTR1A (ABY85410.1), Hs5HTR7 (P34969), Rn5HTR7 (P32305), Pr5HTR7 (AMQ67549.1), Am5HTR7 (NP_001071289.1), Dm5HTR7 (NP_524599.1), Ae5HTR7 (Q9GQ54), Hs5HTR4 (Q13639), Cp5HTR4 (O70528), Rn5HTR4 (Q62758), Mm5HTR4 (P97288), Hs5HTR6 (P50406), Pt5HTR6 (Q5IS65), It5HTR6 (XP_005317590.1), Rn5HTR6 (P31388), Mm5HTR6 (Q9R1C8), Cp5HTR6 (XP_003471412.1), Pm5HTR2A (KPJ17794.1), Pr5HTR2A (XP_022112310.1), Bm5HTR2A (NP_001296483.1), Cl5HTR2A (XP_014254278.1), Rp5HTr2B (AKQ13312.1), Mq5HTR2A (KOX78271.1), Am5HTr2B (NP_001189389.1), Hs5HTR2A (NP_000612.1), Hs5HTR2C (NP_000859.1), Hs5HTR2B (NP_000858.3), Pt5HTR2C (XP_015921531.1), Mp5HTR2C (XP_022169104.1), Am5HTr2A (NP_001191178.1), Pr5HTR2C (XP_022122944.1), Ba5HTR2C (XP_023955125.1), Rn5HTR5B (P35365), Mm5HTR5B (P31387), Hs5HTR5A (NP_076917.1), Rn5HTR5A (P35364), Mm5HTR5A (P30966), Hs5HTR3C (NP_570126.2), Hs5HTR3E (NP_001243542.1), Hs5HTR3D (NP_001157118.1), Hs5HTR3B (NP_006019.1), Rn5HTR3B (NP_071525.1), Hs5HTR3A (AP35868.1), Cp5HTR3A (O70212), Rn5HTR3A (NP_077370.2), Mm5HTR3A (P23979). The phylogenetic tree was constructed using neighbor-joining method. Bootstrap values on branch nodes were obtained after 1,000 repetitions. **(B)** Amino acid sequence alignment of Se-5HTR with orthologous receptors from *Homo sapiens* (Hs-5HTR7: P34969), *Drosophila melanogaster* (Dm-5HTR7: NP_524599.1), *Manduca sexta* (Ms-5HTR7: AGL46976.1), and *Pieris rapae* (Pr-5HTR7: AMQ67549.1). Identical residues among these five sequences are illustrated as white letters against black. Dashes within sequences indicate gaps introduced to maximize homology. Putative seven transmembrane domains (TM1-TM7) are shown as blue bars. Potential N-glycosylation sites 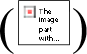, potential phosphorylation sites for protein kinase A and/or C 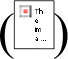, potential residue to interacts with 5-HT amino group 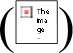 and 5-HT hydroxyl group 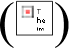, potential disulfide bond groups 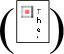, and potential post translational palmitoylation sites 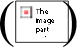 are indicated. Overbars indicate unique motif to aminergic receptor, agonist mediated sequestration and resensitization motif, and PDZ-domain binding motif, respectively. Conserved domains were determined using InterPro tool (https://www.ebi.ac.uk/interpro/) and Prosite (http://prosite.expasy.org/) whereas other residues and motifs were predicted using several tools from DTU bioinformatics (www.cbs.dtu.dk/services/).

*Se-5HTR* sequence attains highly conserved amino acid residues (Fig 1B) found uniquely in the biogenic monoamine receptor family [49–51]. Negatively charged Asp residue in TM3 (Asp^163^) might interact with positively charged amino group of the ligand. A hydrogen bond between the hydroxyl group of serine residue in TM5 (Ser^247^) and the hydroxyl group of 5-HT was predicted. In TM6, the consensus sequence unique to aminergic receptors (Phe^444^-X-X-X-Trp^448^-X-Pro^450^-X-Phe^452^) is conserved. Like other GPCRs, a possible motif (Asn^485^-Pro^486^-X-X-Tyr^489^) that might participate in agonist-mediated sequestration and re-sensitization of Se-5HTR is conserved in TM7. The C-terminus of the receptor contains two potential post-translational palmitoylation cysteine residues (Cys^508^ and Cys-^532^) and PDZ (<u>p</u>ost synaptic density protein, *<u>D</u>rosophila* disc large tumor suppressor, and <u>z</u>onula occludens-1 protein)-domain binding motif (Glu^561^-Ser^562^-Phe^563^-Leu^564^).

### Se-5HTR is expressed in all developmental stages and larval tissues

Expression of *Se-5HTR* in different developmental stages of *S. exigua* was assessed by RT-PCR. Results showed its expression from egg to adult stages (Fig 2A). RT-qPCR revealed variation in its expression among developmental stages, with L5 larvae and adults showing the highest expression levels. In L5 larvae, all tissues analyzed by RT-PCR showed its expression (Fig 2B). RT-qPCR revealed that the midgut exhibited the highest expression level of *Se-5HTR*. Hemocytes and brain also showed relatively high levels of its expression. Basal expression levels of *Se-5HTR* were highly up-regulated in response to immune challenge (Fig 2C). *Se-5HTR* expression was increased ~125-fold after challenge with fungus, *Beauveria bassiana* compared to control (unchallenged). It was increased 60~90-fold after bacterial challenge compared to naïve larvae.

**Fig. 2.**
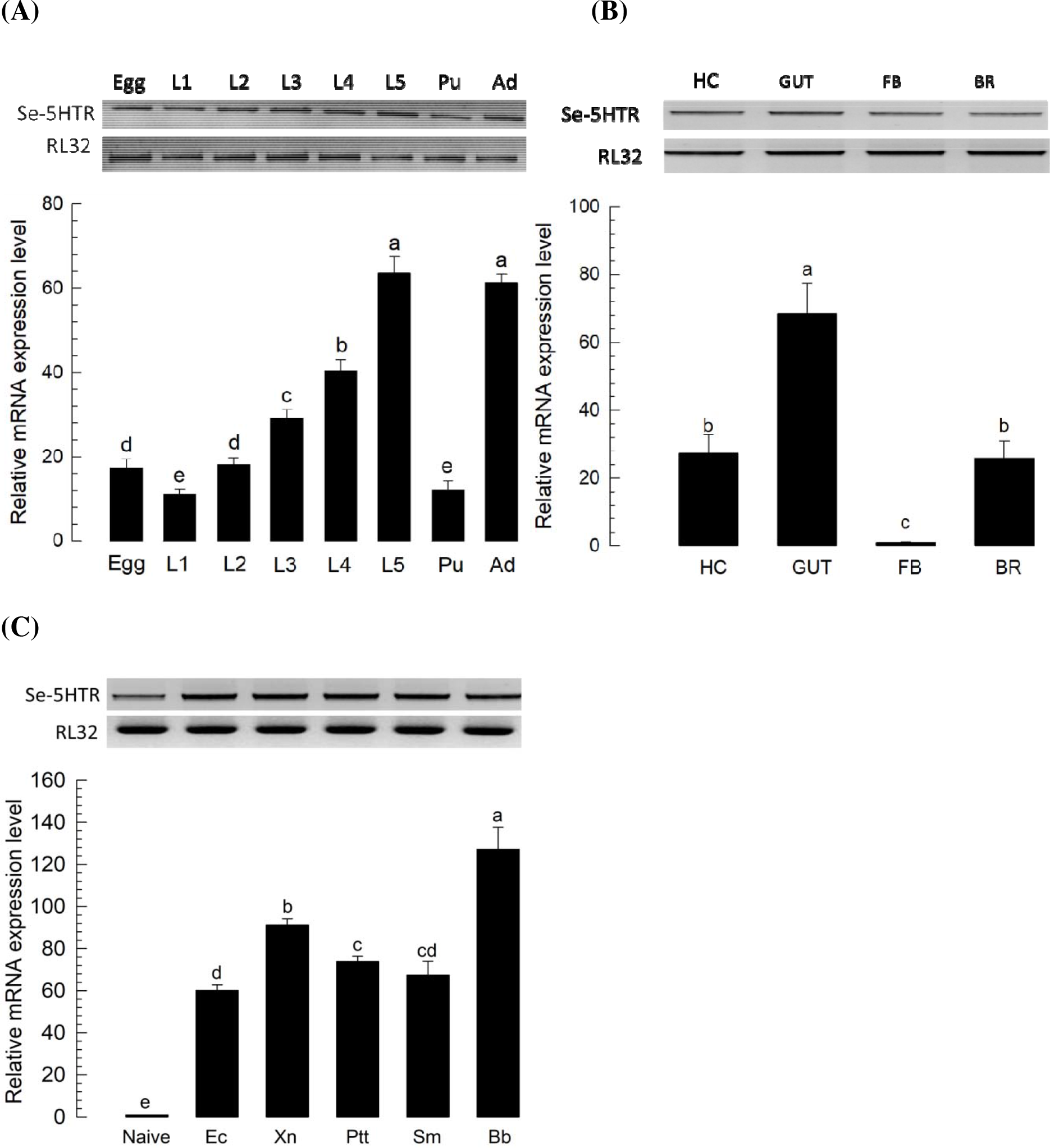
Expression profile of *Se-5HTR.* **(A)** Differential expression of *Se-5HTR* in different developmental stages: egg, larval instars (‘L1-L5’), pupa (‘Pu’), and adult (‘Ad’). **(B)** Differential expression of *Se-5HTR* in different tissues of L5 larvae: hemocyte (‘HC), midgut (‘GUT’), fat body (‘FB’), and brain (‘BR’). **(C)** Expression pattern of *Se-5HTR* in immune-challenged L5 larvae: *Escherichia coli* (‘Ec’), *Xenorhabdus nematophila* (‘Xn’), *Photorhabdus temperata temperata* (‘Ptt’), *Serratia marcescens* (‘Sm’), and *Beauveria bassiana* (‘Bb’). Immune challenge was performed by injecting each L5 larva with 1.8 × 10^5^ cells of bacteria or 5 × 10^5^ conidia of fungi. After incubation at 25°C for 8 h, gene expression analysis was performed using RT-PCR and RT-qPCR. As a constitutive expressional control, a ribosomal gene, *RL32,* was used for expression analysis in RT-PCR and RT-qPCR. Each measurement was replicated three times with independent biological samples. Histogram bars annotated with the same letter are not significantly different at Type I error = 0.05 (LSD test).

### Specific inhibitors against 5-HTR suppress hemocyte behaviors of *S. exigua*

A commercial inhibitor (SB-269970) specific to 5-HT_7_ was used to assess its effect on hemocyte behaviors of *S. exigua* (Fig 3). Total hemocyte count (THC) of L5 larvae was ~1.2 × 10^7^ cells/mL (Fig 3A). THC was significantly (*P* < 0.05) increased in response to 5-HT or bacterial challenge. SB-269970 prevented the increase of THC in response to bacterial challenge. Its IC_50_ value was estimated to be 1.925 μM. These results suggest that Se-5HTR can mediate hemocyte mobilization in response to 5-HT upon bacterial challenge.

**Fig. 3.**
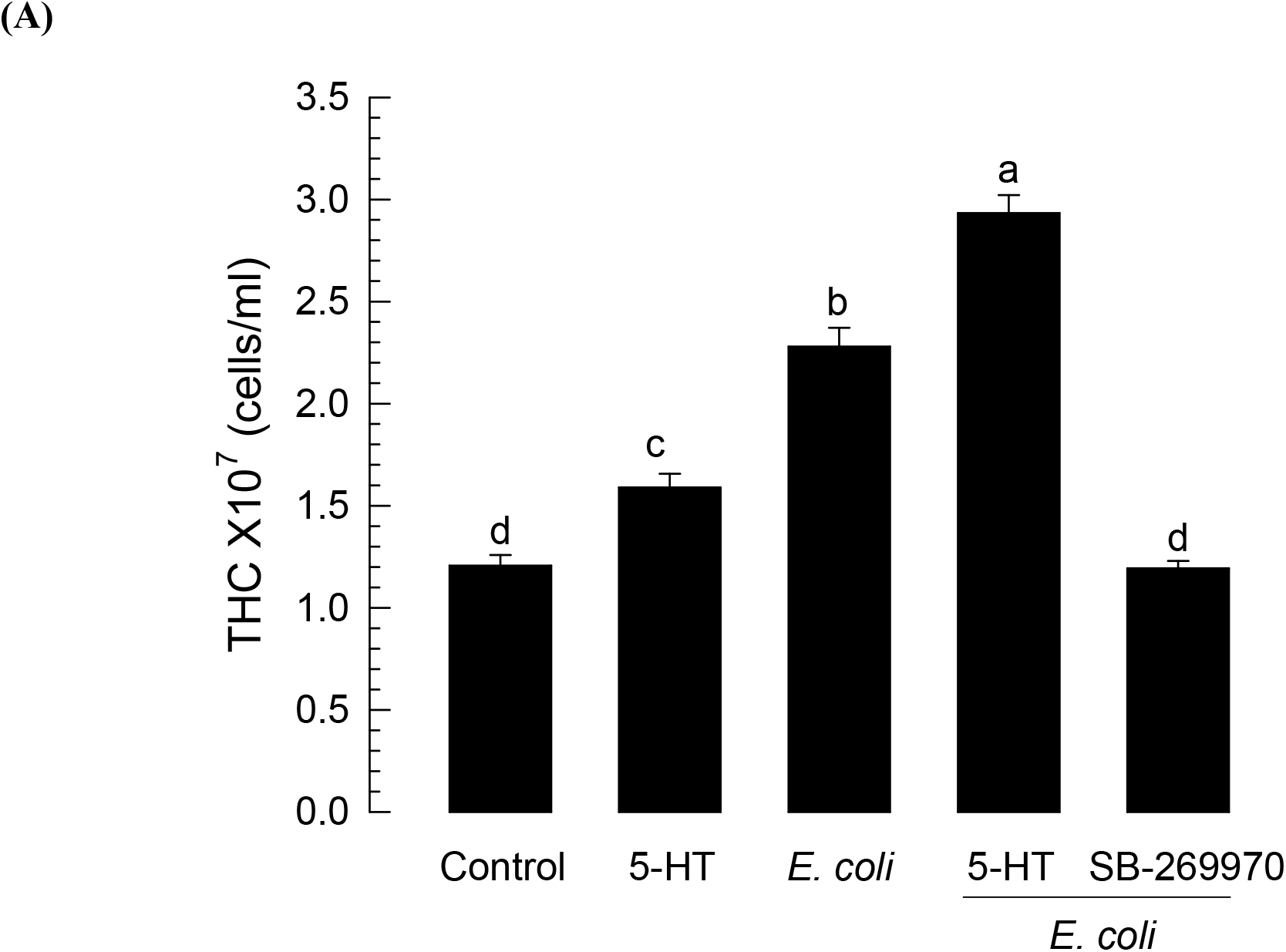
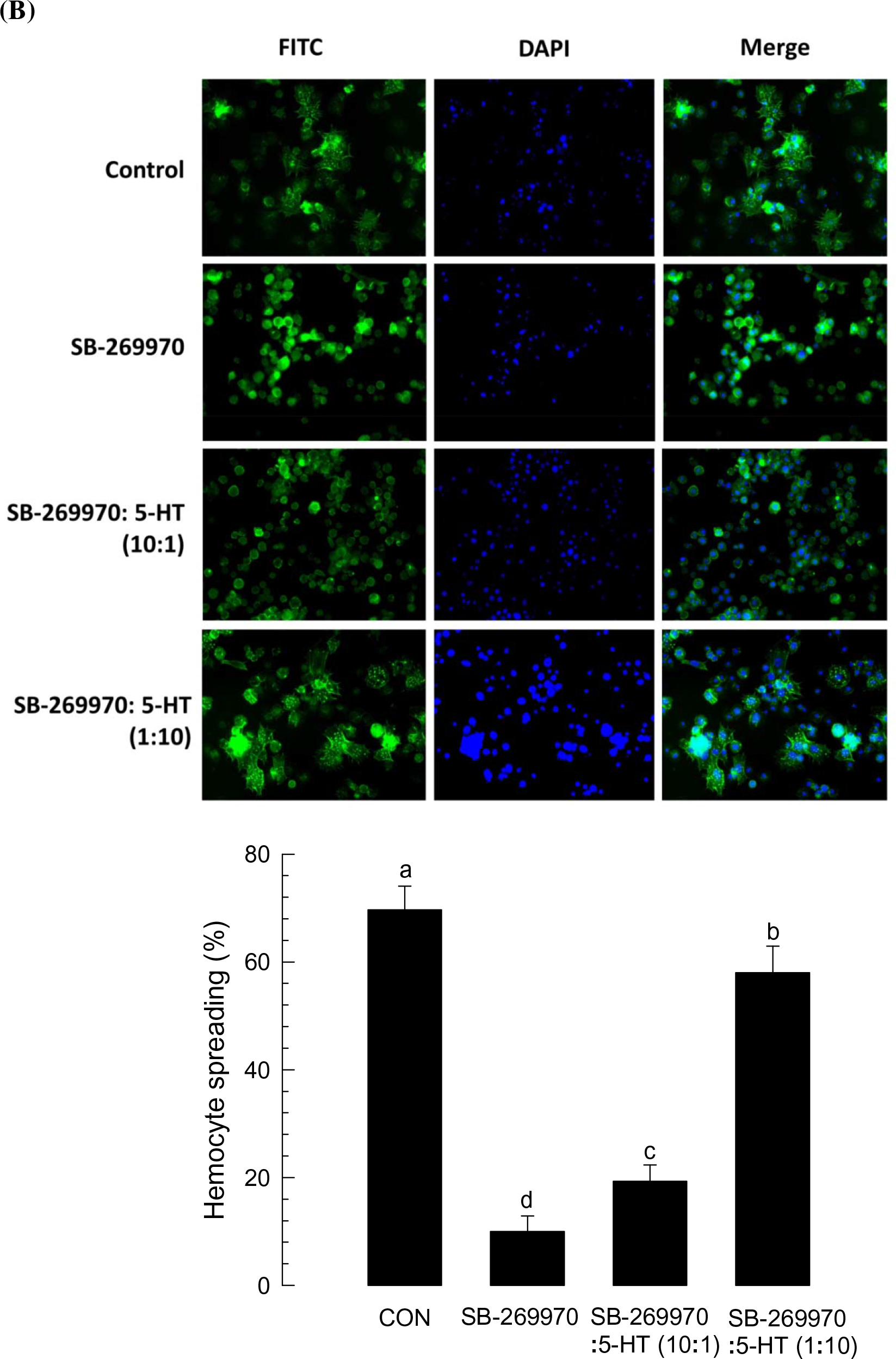
Effect of an inhibitor specific to 5-HT_7_ receptor (‘SB-269970’) on hemocyte behaviors of *S. exigua.* **(A)** Total hemocyte count (THC) analysis in L5 larvae. For the assay, 2 μg of 5-HT or 2 μg of SB-269970 was injected. THC was assessed with or without bacterial coinjection (1.8 × 10^5^ cells per larva). **(B)** Hemocyte-spreading behavior analysis. Hemocyte-spreading was assessed with a 20 μL reaction mixture containing 1 or 2 μL test chemical with 18 or 19 μL hemocytes. 5-HT was co-applied with SB-269970 at ratios of 1:10 (2 μg 5-HT: 20 μg SB-269970) and 10:1 (20 μg 5-HT: 2 μg SB-269970). Spread cells were stained with F-actin and FITC-labeled phalloidin. Nuclei were stained with DAPI. Each treatment was independently replicated three times. Histogram bars indicate percentages of spread hemocytes and error bars indicate standard deviation. Histogram bars annotated with the same letter are not significantly different at Type I error = 0.05 (LSD test).

To determine the modulation effect of Se-5HTR on hemocyte-spreading behavior, competitive inhibition between SB-269970 and 5-HT against Se-5HTR was assessed (Fig 3B). Hemocytes were spread on slide glass with growth of F-actin. However, SB-269970 significantly (*P* < 0.05) suppressed such hemocyte-spreading behavior. The inhibitory effect of the inhibitor was rescued by addition of 5-HT. A relatively low dose (SB-269970: 5-HT = 10:1) of 5-HT did not rescue the inhibitory effect. However, a relatively high dose (1:10) of 5-HT significantly (*P* < 0.05) rescued such inhibitory effect, indicating a specific competition between inhibitor and ligand against Se-5HTR.

### RNA interference (RNAi) of Se-5HTR

To address the immunomodulatory effect of Se-5HTR on hemocytes, its gene expression was knocked-down by RNAi (Fig 4). RNAi was performed using dsRNA specific to *Se-5HTR.* dsRNA-injected larvae exhibited significant (*P* < 0.05) reduction of *Se-5HTR* expression. RT-qPCR analysis showed that about 90% of *Se-5HTR* mRNA was suppressed by dsRNA treatment at 24 h PI (Fig 4A). At 24 h PI of dsRNA, *Se-5HTR* expression was analyzed in four different tissue samples. Compared to control tissues, dsRNA-treated larvae exhibited significant (*P* < 0. 05) reduction of *Se-5HTR* expression in all tissues including hemocytes.

**Fig. 4.**
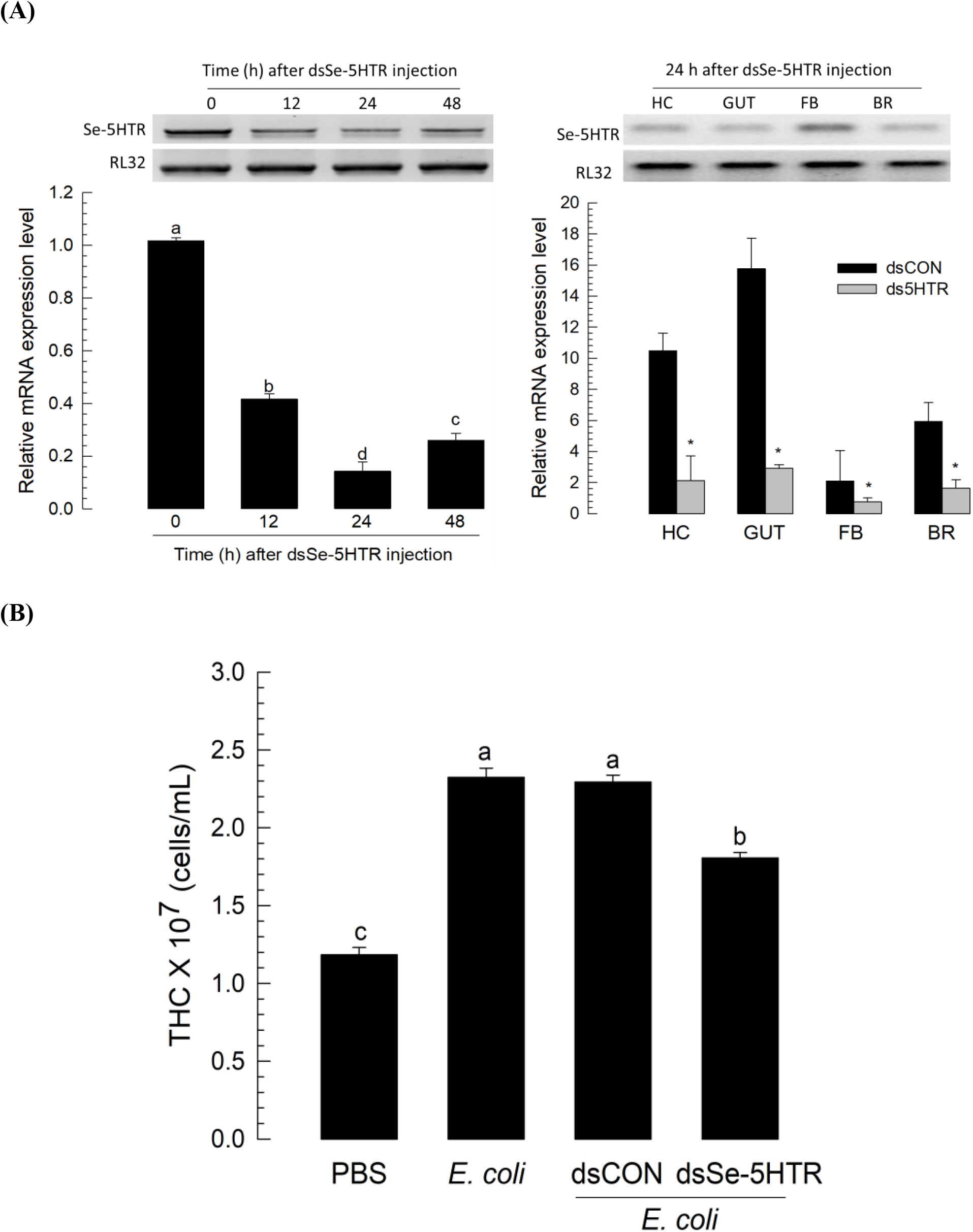
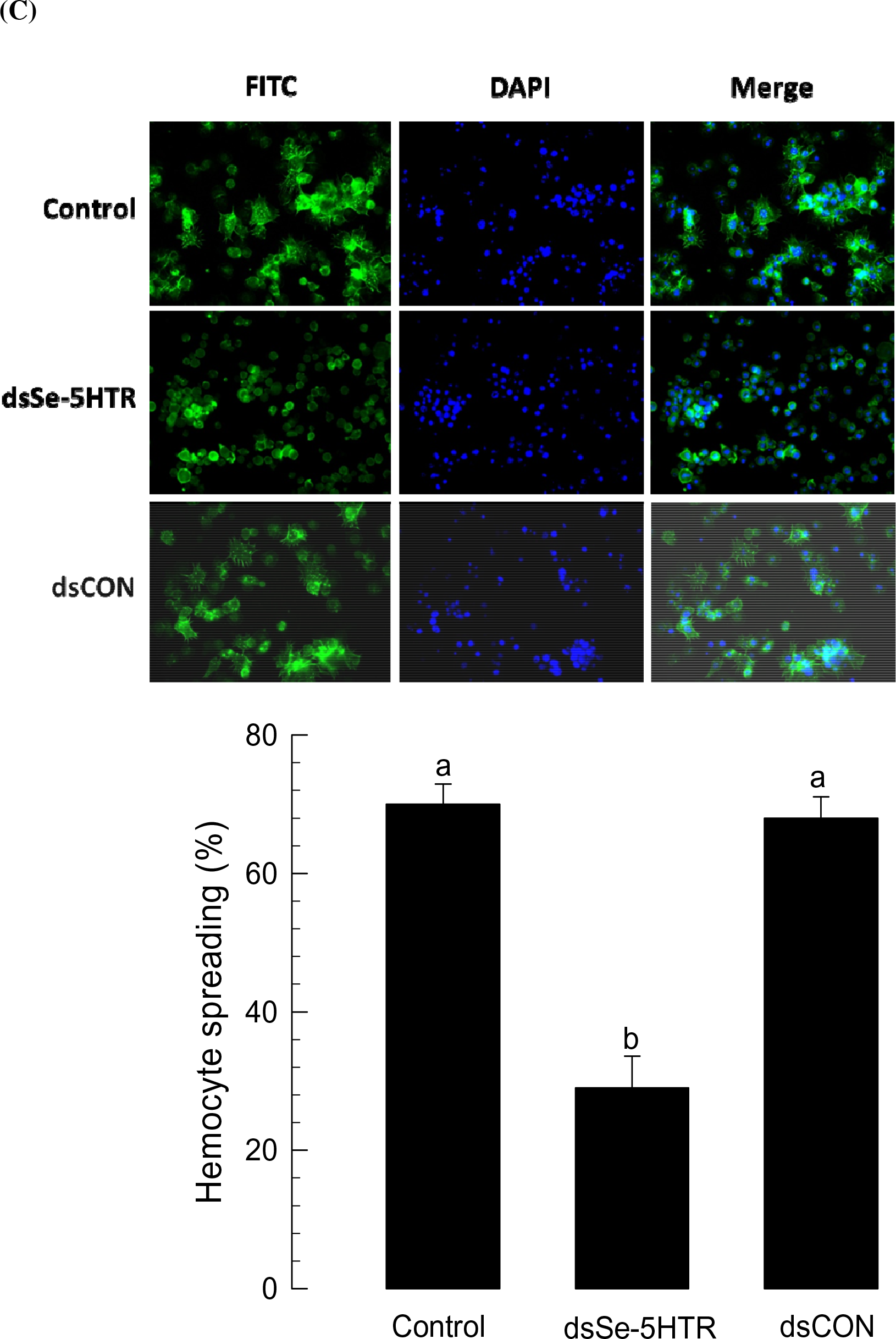
RNA interference (RNAi) of *Se-5HTR* and suppression of hemocyte behaviors. **(A)** RNAi of *Se-5HTR* expression with a gene-specific dsRNA (dsSe-5HTR). About 900 ng of dsSe-5HTR was injected to L5. At 0, 12, 24, and 48 h post-injection (PI), expression levels of *Se-5HTR* were assessed from whole body. Expression levels of *Se-5HTR* in different tissue parts were assessed at 24 h PI. As a constitutive expressional control, a ribosomal gene, *RL32,* was used for expression analysis in RT-PCR and RT-qPCR. As a control RNAi (dsCON), dsRNA specific to *CpBV-ORF302* (a viral gene) was used for expression analysis. In RT-qPCR, each treatment was triplicated. **(B)** Influence of RNAi on up-regulation of total hemocyte count (THC) in response to bacterial challenge. At 24 h PI of dsSe-5HTR, hemolymph was collected from L5 larvae for THC assessment. Each treatment was replicated three times. **(C)** Influence of RNAi on hemocyte-spreading behavior. For spreading assay, hemocytes from larvae treated with dsRNA were collected at 24 h. Each treatment was independently replicated three times. Spread cells were stained with F-actin and FITC-labeled phalloidin. Nuclei were stained with DAPI. Histogram bars indicate percentages of spread hemocytes and error bars indicate standard deviation. Different letters above standard error bars indicate significant difference among means at type I error = 0.05 (LSD test). Asterisks represent significant difference between control and treatment in each tissue.

### RNAi of Se-5HTR suppresses hemocyte behaviors of *S. exigua*

The effect of RNAi specific to Se-5HTR on hemocyte mobilization in response to bacterial challenge was analyzed (Fig 4B). Upon bacterial challenge, THC increased by more than two folds. However, RNAi treatment significantly (*P* < 0.05) suppressed such increase of THC. The RNAi treatment also significantly (*P* < 0.05) influenced hemocyte-spreading behavior (Fig 4C). On glass slide, hemocytes exhibited spreading behavior in 40 min by cytoskeletal rearrangement through F-actin growth (see phalloidin staining). However, hemocytes collected from larvae treated with RNAi specific to Se-5HTR lost such behavior.

### Modulation of cellular immune responses by Se-5HTR

The influence of Se-5HTR on modulating hemocyte-spreading behavior suggested that it might mediate cellular immune responses against bacterial infection. Phagocytosis against FITC-labeled *Escherichia coli* was observed in hemocytes from control larvae. Labeled bacteria were observed within hemocytes of control larvae (Fig 5A). However, hemocytes of larvae treated with dsRNA specific to Se-5HTR significantly (*P* < 0.05) lost such phagocytosis, similar to that found for hemocytes of larvae treated with SB-269970. Phagocytosis was decreased around 57% or 78% after treatment with dsRNA or inhibitor, respectively. In response to bacterial challenge, *S. exigua* formed ~78 hemocytic nodules per larva (Fig 5B). However, RNAi specific to Se-5HTR or inhibitor treatment significantly (*P* < 0.05) reduced such cellular immune response.

**Fig. 5.**
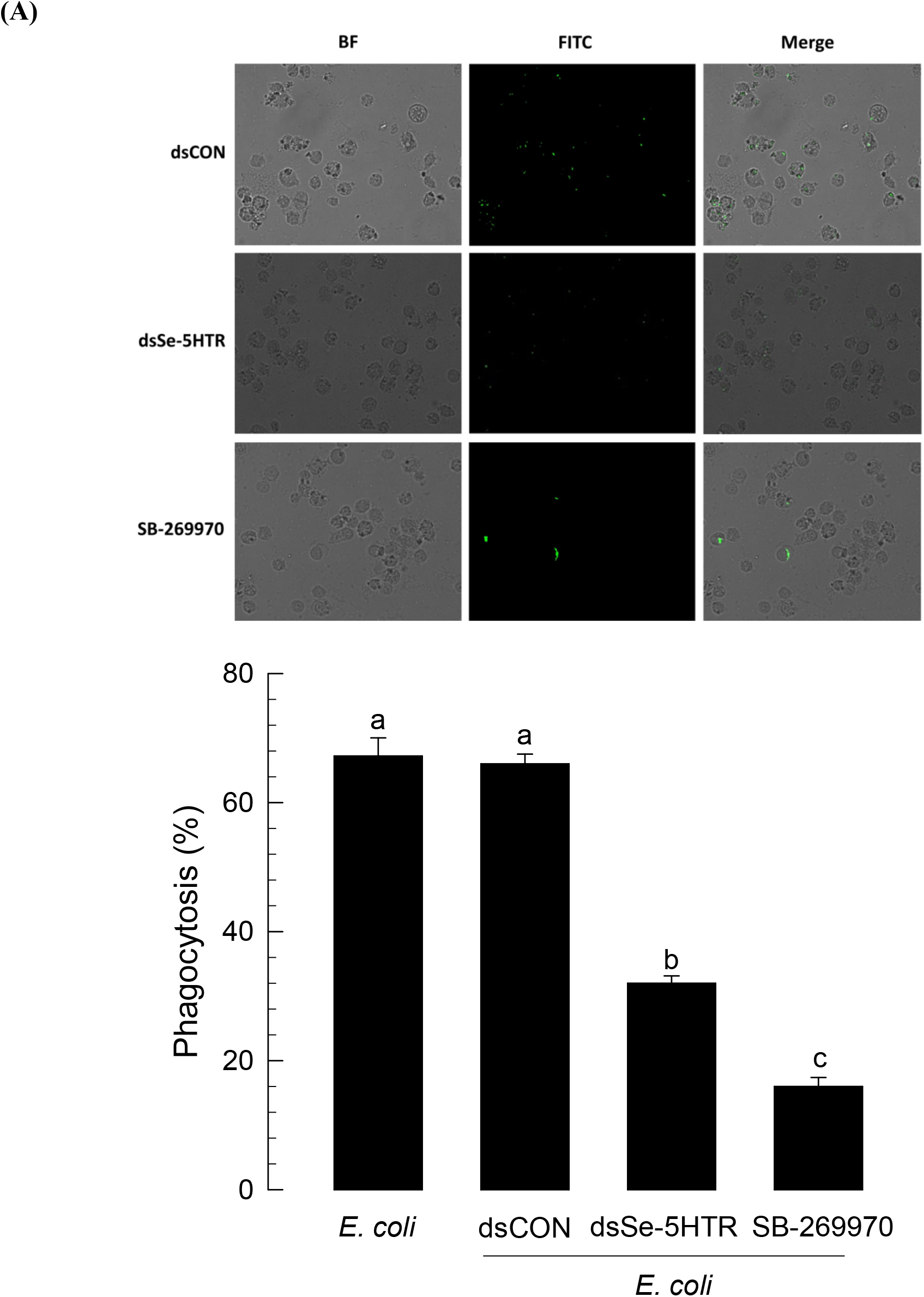
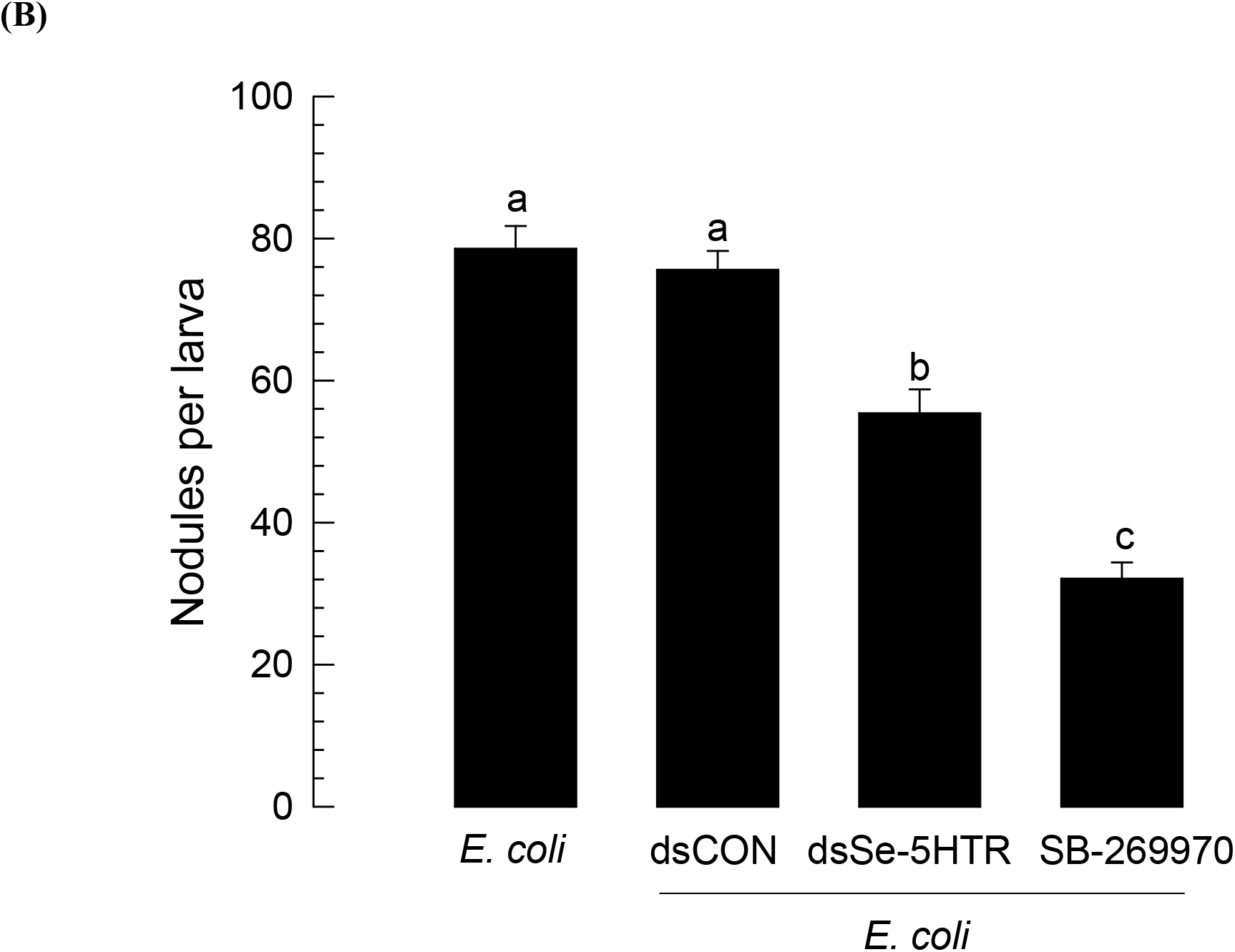
RNA interference (RNAi) of *Se-5HTR* and suppression of cellular immune responses. **(A)** Analysis of phagocytosis in L5 larvae of *S. exigua.* FITC-tagged *E. coli* were injected into L5 larvae at 24 h post-injection of dsSe-5HTR. A specific inhibitor (SB-269970) to 5-HT_7_ receptor was co-injected at a dose of 2 μg along with FITC-tagged bacteria. At 15 min after injection, phagocytic cells were observed under a fluorescent microscope at 400 × magnification and percentage of phagocytotic hemocytes was calculated from randomly chosen 100 cells. Total hemocytes were observed from bright-field (‘BF’). Each treatment was independently replicated three times. **(B)** Nodulation assay. After RNAi, *E. coli* (1.8 × 10^5^ cells/larva) was injected to L5 larvae. SB-269970 (2 μg/larva) was co-injected with bacteria. After 8 h incubation at 25°C, treated insects were assessed for nodule formation. Histogram bars indicate percentages of phagocytosis and error bars indicate standard deviation. Histogram bars annotated with the same letter are not significantly different at Type I error = 0.05 (LSD test).

### Influence of bacterial secondary metabolites on immune responses mediated by Se-5HTR

RNAi or specific inhibitor assays indicated that 5-HTR could mediate both cellular immune responses of phagocytosis and nodule formation in response to bacterial challenge. Bacterial metabolites of two entomopathogens, *Xenorhabdus nematophila* (Xn) and *Photorhabdus temperata temperata* (Ptt), were extracted from their culture broth using different organic solvents and their inhibitory activities against cellular immune responses were then assessed (Fig 6). Organic solvent extracts of Xn- or Ptt-cultured broth exhibited significant (*P* < 0.05) inhibitory activities against phagocytosis (Fig 6A) and nodulation (Fig 6B), although there were variations in their inhibitory activities among extracts.

**Fig. 6.**
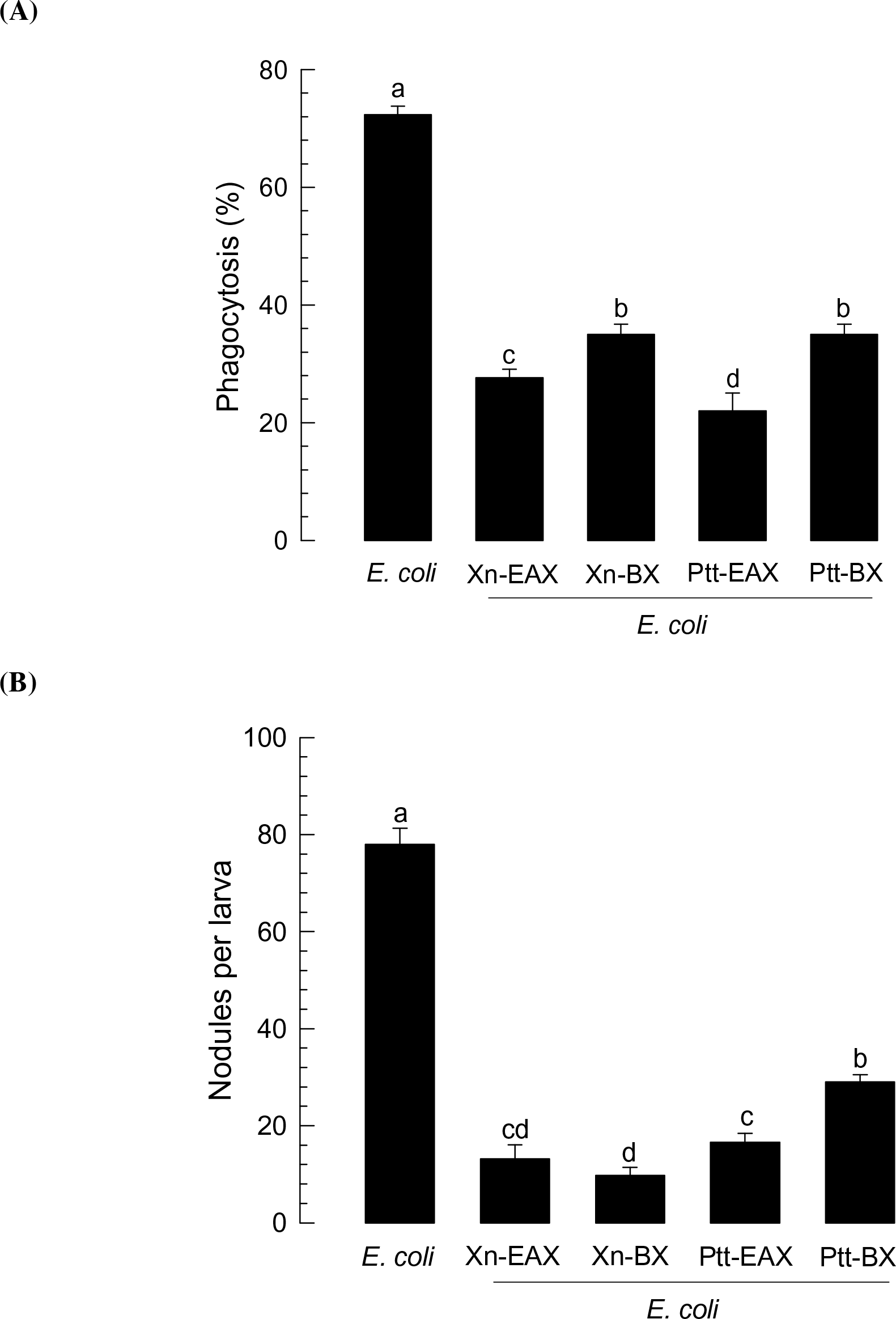
Immunosuppressive activities of organic extracts from culture broth of *Xenorhabdus nematophila* (‘Xn’) and *Photorhabdus temperata temperata* (‘Ptt’) bacteria against L5 larvae of *S. exigua.* Each larva was injected with *E. coli* (10^5^ cells) along with 1 μL of ethyl acetate (‘EAX’) or butanol (‘BX’) extract. **(A)** Phagocytosis in L5 larvae. FITC-tagged *E. coli* were injected to L5 larvae at 24 h post-injection of an organic extract. After 15 min, phagocytic cells were observed under a fluorescent microscope at 400 × magnification and percentage of phagocytotic hemocytes was calculated from randomly chosen 100 cells. Each treatment was independently replicated three times. **(B)** Nodule formation. *E. coli* (1.8 × 10^5^ cells/larva) was injected to L5 larvae. An organic extract was co-injected with the bacteria. After 8 h of incubation at 25°C, treated insects were assessed for nodule formation. Each treatment was replicated with 10 larvae. Different letters above standard deviation bars indicate significant difference among means at Type I error = 0.05 (LSD test).

To identify bacterial secondary compounds that could inhibit Se-5HTR, 37 compounds (HB4 – HB602) derived from *Xenorhabdus* and *Photorhabdus* [47] were screened for their inhibitory activities against hemocyte nodule formation and phagocytosis (Fig 7). More than three 75% (28 out of 37 compounds) of these test compounds exhibited significant (*P* < 0.05) inhibition against phagocytosis (Fig 7A). In nodulation assay, all test compounds exhibited significant (*P* < 0.05) inhibition (Fig 7B). Since Se-5HTR could mediate both cellular immune responses, bacterial compounds that highly inhibited both phagocytosis (< 60%) and nodulation (< 30 nodules per larva) were selected. As a result, 10 potent chemicals (HB 4, HB 5, H23, HB 30, HB44, HB 45, HB 50, HB 223, HB 302, and HB 531) belonging to six chemical categories [phenylethylamide (PEA), tryptamide, xenortide, xenocycloin, nematophin, and GameXPeptide] were found (S2 Fig). All these compounds exhibited median inhibitory concentration (IC_50_) of 58~253 μM against cellular immune response of hemocytic nodulation.

**Fig. 7.**
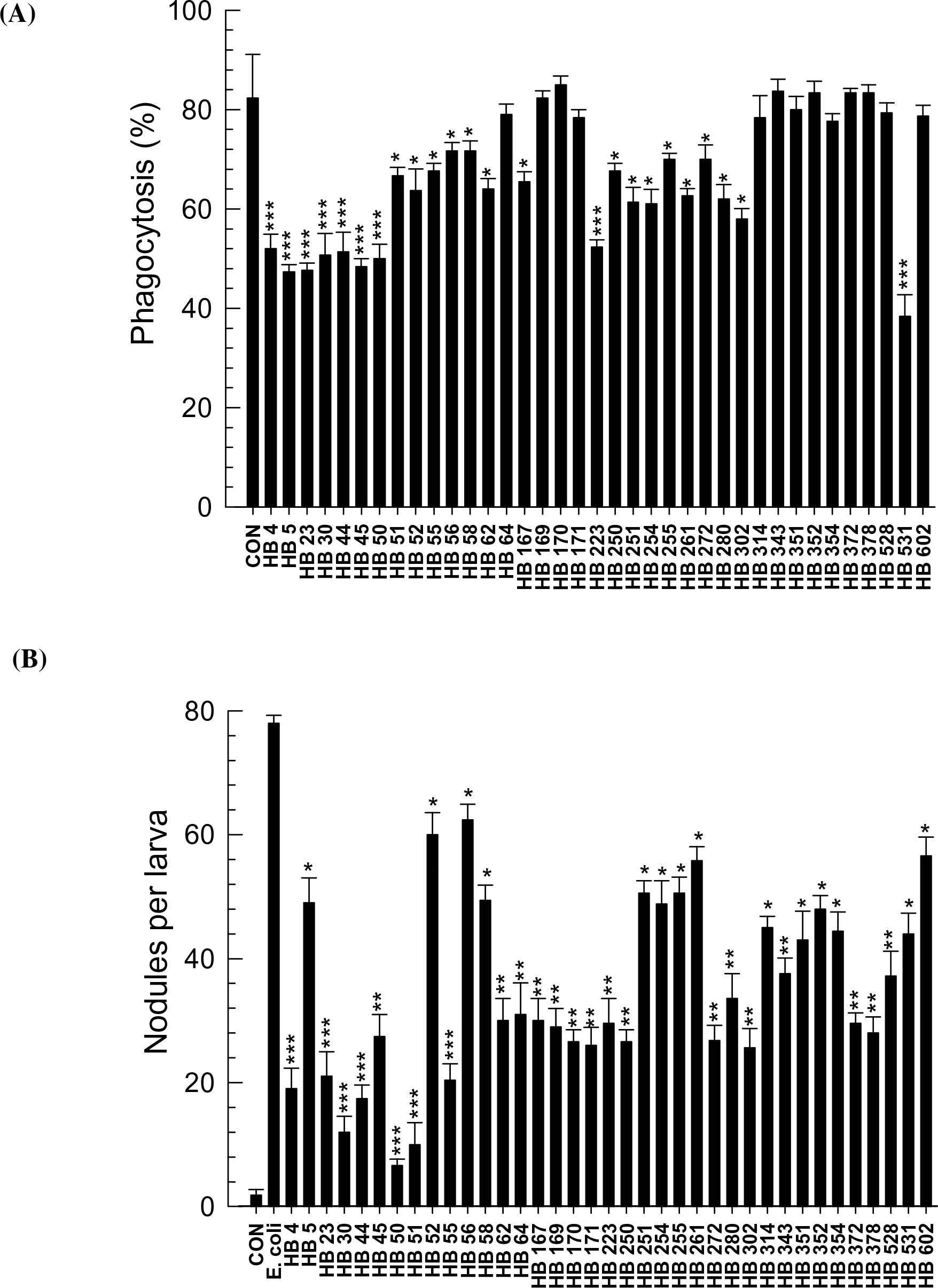
Screening 37 bacterial secondary metabolites derived from *X. nematophila* and *P. temperata temperata* for their effects on cellular immune responses mediated by Se-5HTR in *S. exigua.* **(A)** Screening with phagocytosis assay. FITC-tagged *E. coli* cells were injected to L5 larvae along with 1 μg of each test chemical. After 15 min, phagocytic cells were observed under a fluorescent microscope at 400 × magnification and percentage of phagocytotic hemocytes was calculated from randomly chosen 100 cells. Each treatment was independently replicated three times. **(B)** Screening with nodule formation assay. *E. coli* (1.8 × 10^5^ cells/larva) was injected to L5 larvae along with 1 μg of each test chemical. After 8 h of incubation at 25°C, treated insects were assessed for nodule formation. Each treatment was replicated with 10 larvae. Asterisks above standard deviation bars indicate significant difference compared to control (‘CON’ without inhibitor) at Type I error = 0.05 (*), 0.01 (**), and 0.005 (***) (LSD test).

To support the specific inhibitory activity of these selected compounds against Se-5HTR, competitive assay was performed between test compounds and 5-HT (Fig 8). With a constant concentration of 5-HT, phagocytotic behavior of *S. exigua* hemocytes gradually decreased with increasing amount of test compound (Fig 8A). On the other hand, increase of 5-HT amount gradually decreased the phagocytotic behavior using a constant amount of test compound (Fig 8B).

**Fig. 8.**
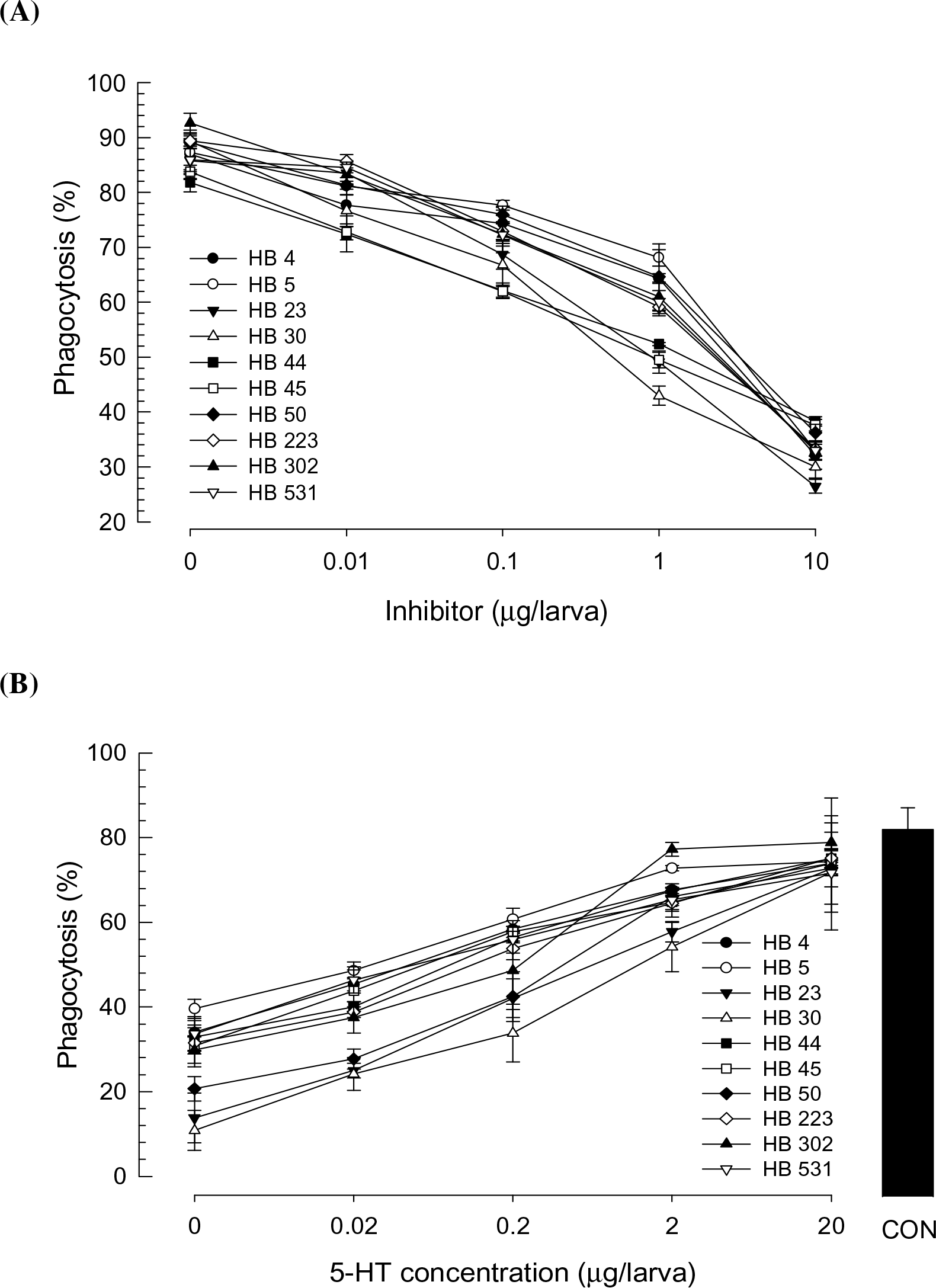
Competitive inhibition of 10 selected bacterial metabolites derived from *X. nematophila* and *P. temperata temperata* with 5-HT against Se-5HTR mediating phagocytosis. FITC-tagged *E. coli* cells were injected to L5 larvae along with test chemicals. After 15 min, phagocytic cells were observed under a fluorescent microscope at 400 × magnification and percentage of phagocytotic hemocytes was calculated from randomly chosen 100 cells. Each treatment was independently replicated three times. **(A)** Dose-response of inhibitory compound (0, 0.01, 0.1, 1, and 10 μg/larva) with a fixed 5-HT concentration (1 μg/larva). **(B)** Dose-response of 5-HT (0, 0.02, 0.2, 2 and 20 μg/larva) with a fixed HB compound concentration (1 μg/larva). ‘CON’ represents a positive control without any inhibitor.

### Influence of PEA derivatives on phagocytosis mediated by Se-5HTR

These 10 selected compounds shared phenyl (or aromatic) ethylamide backbone except xenocycloin (S2 Fig). Based on PEA backbone, 45 derivatives were selected from a chemical bank and their inhibitory activities against the phagocytotic behavior of *S. exigua* hemocytes were tested (Fig 9A). More than 66% (30 out of 45 compounds) of PEA derivatives exhibited significant (*P* < 0.05) inhibition against phagocytosis. Three PEA derivatives (Ph15, Ph17, Ph33) had inhibitory activities similar to a bacterial metabolite (HB 44), with IC_50_ at 1.8 ~ 5.6 μM against phagocytosis (Fig 9B). These three PEA derivatives competitively inhibited the phagocytosis mediated by Se-5HTR (Fig 9C).

**Fig. 9.**
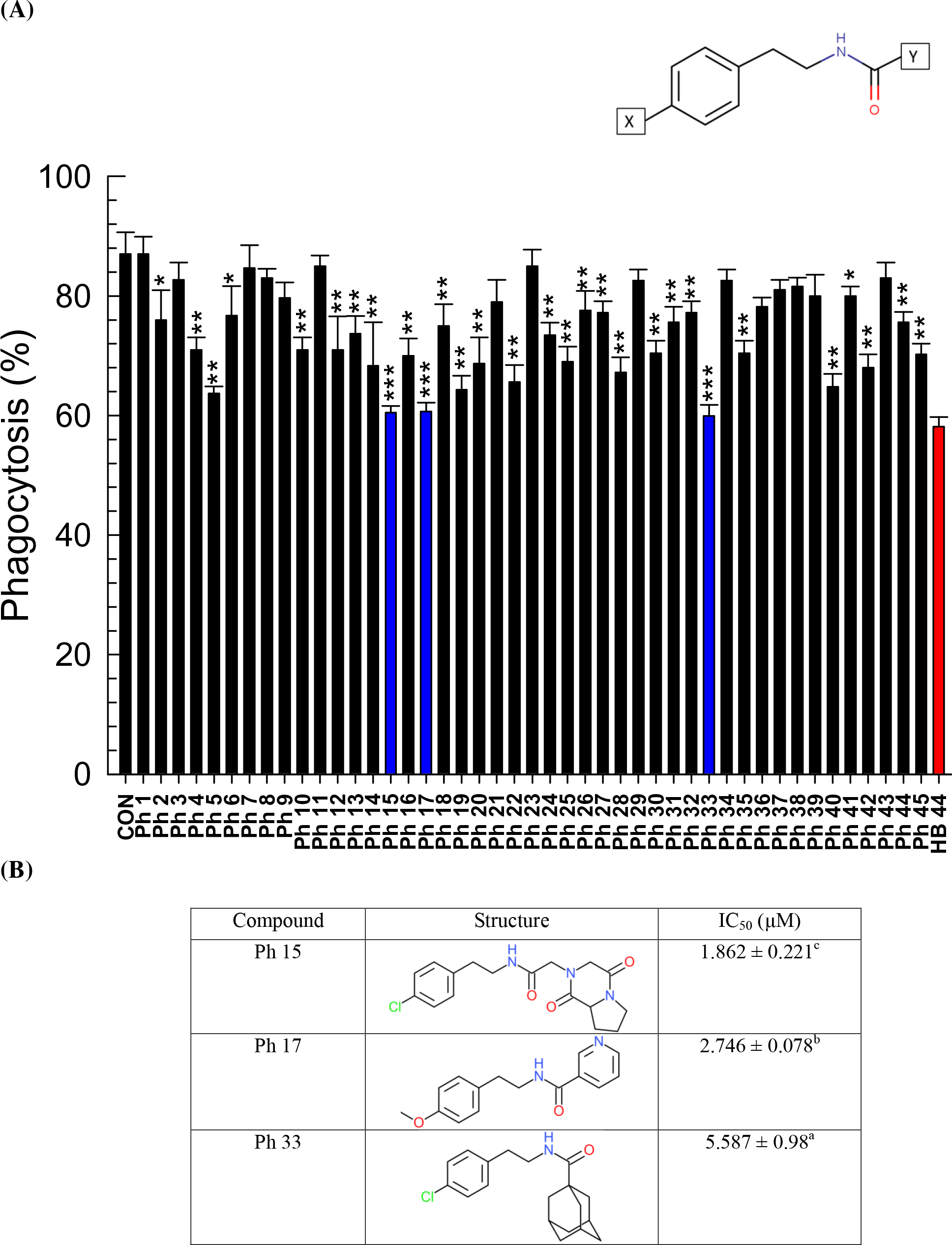
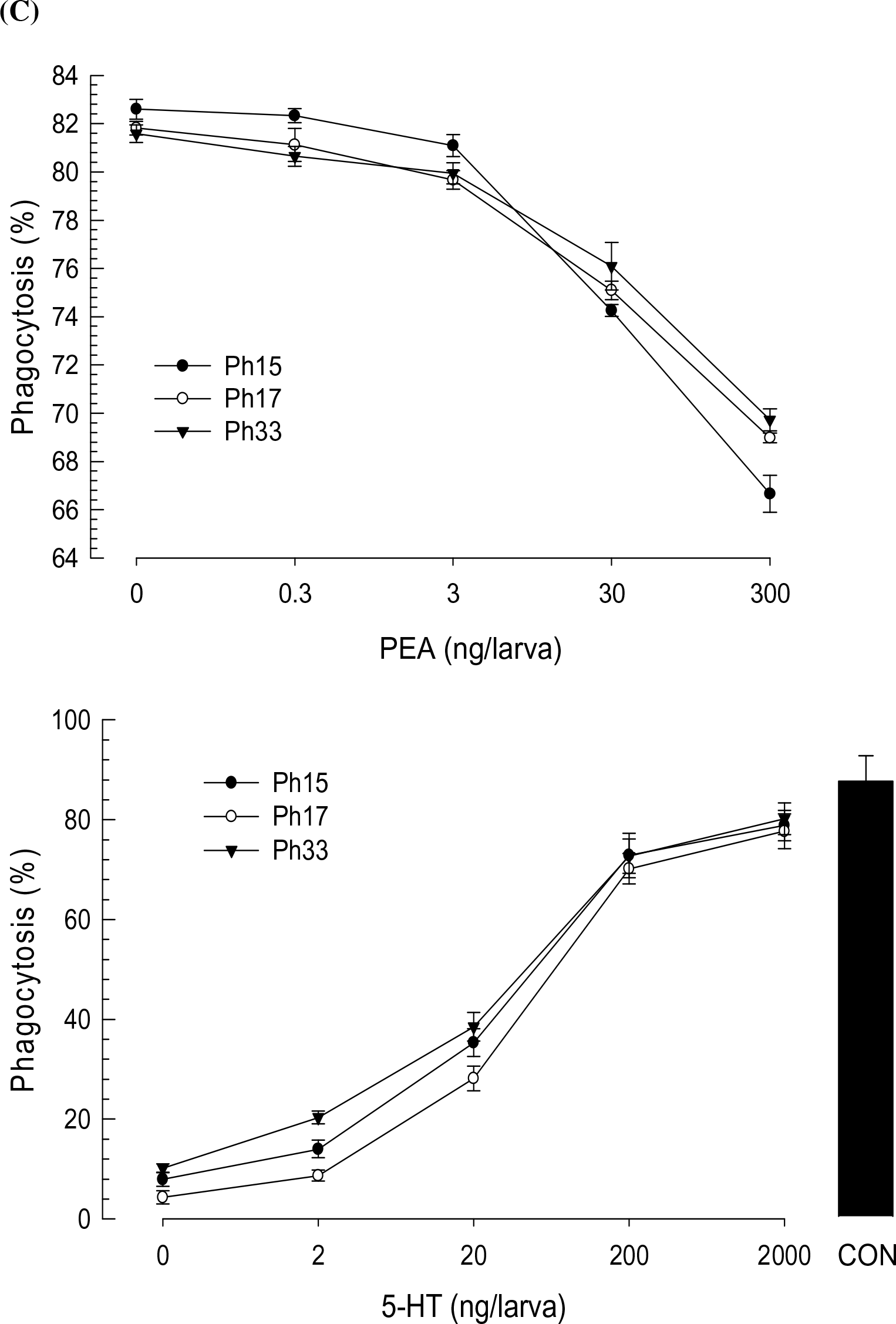
Validation of phenylethylamide (PEA) compounds for their inhibitory activities against 5-HTR using derivatives with different side chains (‘X’ and ‘Y’ of PEA skeleton). **(A)** Screening of 45 PEA derivatives for their effects on hemocyte phagocytosis of *S. exigua* with HB 44, a potent bacterial metabolite, as reference. PEA compounds (300 ng/larva) were injected into hemocoels of L5 larvae along with FITC-tagged bacteria. After incubating for 15 min, hemocytes were assessed for phagocytosis. Asterisks above standard deviation bars indicate significant difference compared to control (‘CON’ without inhibitor) at Type I error = 0.05 (*), 0.01 (**), and 0.005 (***) (LSD test). **(B)** Chemical structures of three selected PEA compounds and their median inhibitory doses (IC_50_). **(C)** Competitive inhibition of the three selected PEA derivatives with 5-HT against Se-5HTR mediated phagocytosis. For dose-response analysis of inhibitory compounds (0, 0.3, 3, 30, and 300 ng/larva), a fixed 5-HT concentration (1 μg/larva) was used. For dose-response of 5-HT (0, 2, 20, 200 and 2,000 ng/larva), a fixed HB compound concentration (1 μg/larva) was used. ‘CON’ represents a positive control without any inhibitor.

### Synthesis of a potent inhibitor and inhibitory efficacy against Se-5HTR

Potent compounds from 45 PEA derivatives were analyzed for their structures and activities against phagocytosis mediated by Se-5HTR (S3 Fig). They shared the PEA backbone. However, they had different side chains at ‘X’ and ‘Y’ (Fig S6A). When different X substituents were compared for their inhibitory activities, methoxy was the most potent moiety (Fig S6B). When different Y substituents were compared for their inhibitory activities with respect to identical X groups, hexahydropyrrolo[1,2-a]pyrazine-1,4-dione was the most potent moiety (Fig S6C). This analysis allowed us to design a hypothetical compound containing methoxy in X and hexahydropyrrolo[1,2-a]pyrazine-1,4-dione at Y based on PEA backbone (Fig. S6D).

The designed compound (‘PhX’) was chemically synthesized and in its inhibitory activity against phagocytosis was assessed compared with a specific 5-HT_7_ inhibitor, SB269970, as a reference (Fig 10). PhX was highly potent. It competitively inhibited cellular immune response with 5-HT. Its inhibitory activity was more potent (*t* = 14.9; df = 32; *P* ≤ 0.0001) than SB269970.

**Fig. 10.**
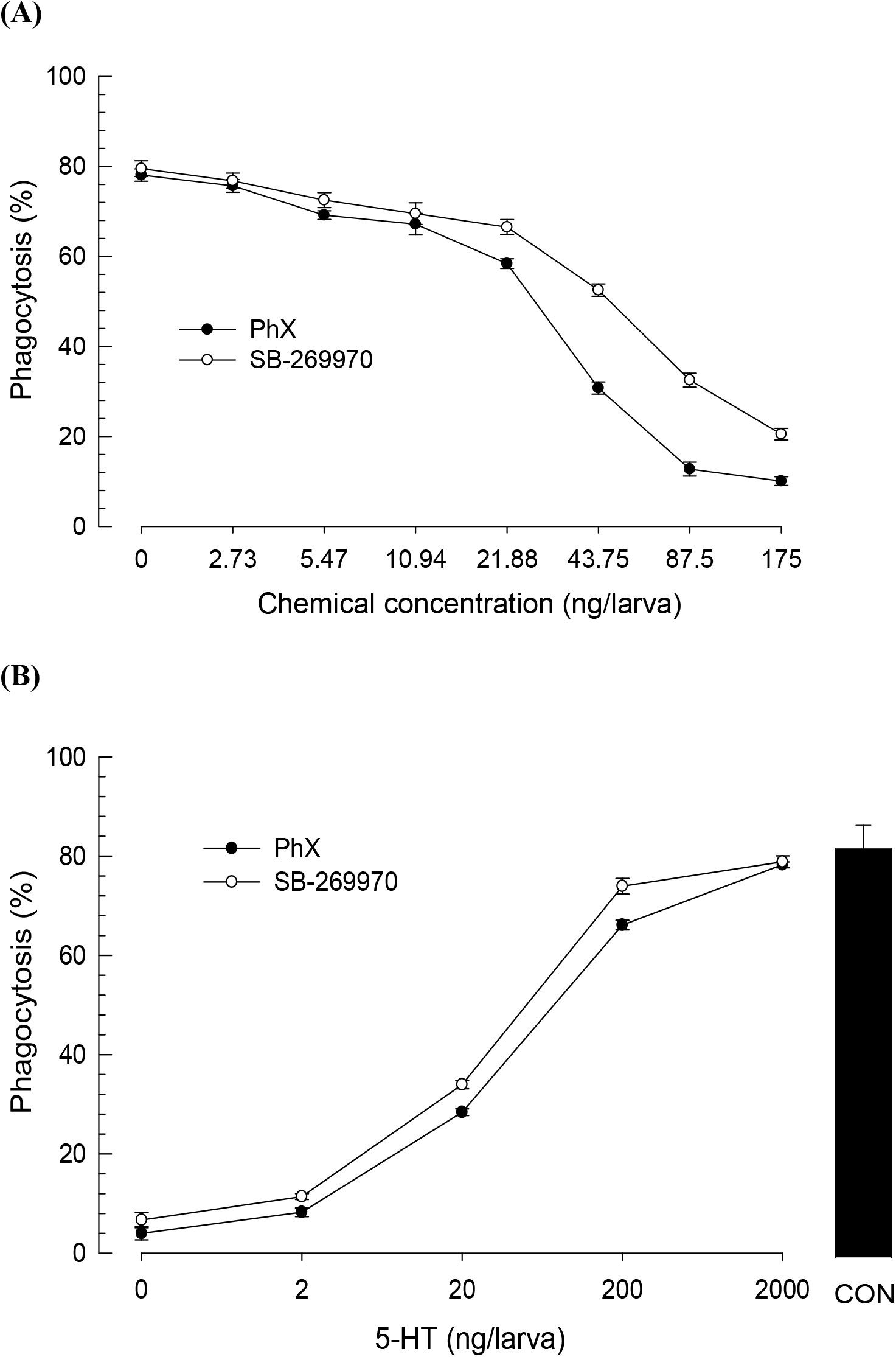
Specific inhibition of a designed compound (PhX) on Se-5HTR with a commercial 5-HTR_7_ inhibitor (SB-269970) as reference in phagocytosis of *S. exigua.* FITC-tagged *E. coli* cells were injected to L5 larvae along with test chemicals. After 15 min, phagocytic cells were observed under a fluorescent microscope at 400 × magnification and percentage of phagocytotic hemocytes was calculated from randomly chosen 100 cells. Each treatment was independently replicated three times. **(A)** Dose-response of PhX and SB-269970 with a fixed 5-HT concentration (500 ng/larva). **(B)** Dose-response of 5-HT (0, 0.002, 0.02, 0.2, and 2 μg/larva) with a fixed inhibitor concentration (10 ng/larva). ‘CON’ represents a positive control without any inhibitor.

## Discussion

5-HT plays a crucial role in mediating cellular immune responses of *S. exigua* by stimulating actin rearrangement [45]. Furthermore, it performs a functional cross-talk with eicosanoid signaling via a small G protein Rac1 [52]. However, its signaling pathway leading to immune responses remains unclear due to the lack of its receptor information in *S. exigua.* This study identified a 5-HT receptor and showed its physiological function in mediating immune responses.

Se-5HTR attains seven transmembrane domains and shares common molecular characters with other 5-HT receptors. Like other GPCRs, Se-5HTR contains the canonical seven transmembrane domains along with consensus glycosylation in the N-terminus (Asn^48^ and Asn^53^) [53]. Additionally, its sequence contains a consensus aspartic acid residue in TM3 (Asp^163^) and a serine residue in TM5 (Ser^247^) to interact with functional groups of biogenic monoamines [54]. At the intracellular border of TM3, the highly conserved Asp^180^-Arg^181^-Tyr^182^ motif is evident for a strong ionic interaction with Glu^430^ residue adjacent to the intracellular end of TM6 that plays a crucial role in GPCR signal transduction [55]. The sequence contains consensus Phe^444^-X-X-X-Trp^448^-X-Pro^450^-X-Phe^452^ motif in TM6 which is unique to biogenic monoamine GPCRs [38]. Additionally, Se-HTR contains two potential post-translational palmitoylation cysteine residues (Cys^508^ and Cys^532^) [56] at the final intracellular region with a PDZ-domain binding motif (Glu^561^-Ser^562^-Phe^563^-Leu^564^) at the C-terminus [57].

*Se-5HTR* is expressed in all developmental stages. It is expressed in immune-associated tissues (hemocytes and fat body), digestive (gut) tissues, and nervous (brain) tissues at larval stage. Three 5-HT receptors of *P. rapae* larvae are all expressed, although their expression levels in tissues are different. Subtypes 1A and 1B receptor are highly expressed in nervous tissues while subtype 7 receptor is mainly expressed in digestive tissue [36]. *Se-5HTR* was also highly expressed in the gut like 5-HT_7_ of *P. rapae.* Furthermore, our phylogenetic analysis of 5-HT receptors showed that these two insect 5-HT_7_s were closely related and clustered. Expression levels of *Se-5HTR* in hemocytes were similar to those in the brain. This suggests that Se-5HTR is associated with immune function as well as neurophysiological function. Indeed, bacterial or fungal infection up-regulated the expression of *Se-5HTR.* The increase of *Se-5HTR* expression might be explained by the up-regulation of *de novo* biosynthesis of its ligand, 5-HT, via increase in expression of biosynthetic genes as seen in hemocytes of *P. rapae* larvae after immune challenge [22].

The presence of 5-HT_7_ receptor in hemocytes of *S. exigua* and its physiological function associated with immune responses were supported by its sensitivity to specific 5-HTR inhibitors. In response to 5-HT or bacterial challenge, *S. exigua* larvae exhibited significant increase of THC. This up-regulation of THC was explained by mobilization of sessile hemocytes to circulatory form by cytoskeletal rearrangement via a small G protein, Rac1 [45,52]. This suggests that Se-5HTR can activate Rac1 to stimulate the hemocyte behavior. In vertebrates, activation of 5-HT_7_ receptor increases cAMP level via a trimeric G protein G_s_ and small G proteins of Rho family including Cdc42, RhoA, and Rac1 via another trimeric G protein G_12_ [58]. This suggests that immune challenge can induce biosynthesis and release of 5-HT which then binds to Se-5HTR on hemocytes and activates Rac1 to stimulate hemocyte behaviors. In addition, the cAMP pathway triggered by Se-5HTR might activate Akt and ERK1/2 as seen in mammalian cancer cells [59] to facilitate actin rearrangement to form cellular shape change of hemocytes. These findings suggest that Se-5HTR plays a crucial role in hemocyte migration and cell shape change during cellular immune responses. Indeed, RNAi of *Se-5HTR* expression resulted in significant immunosuppression by exhibiting reduction in phagocytosis and nodulation.

Bacterial metabolites derived from two entomopathogens, *Xenorhabdus* and *Photorhabdus,* inhibited cellular immune responses mediated by Se-5HTR. Especially, six chemical groups (phenylethylamide, tryptamide, xenortide, xenocycloin, nematophin, and GameXPeptide) highly inhibited both phagocytosis and nodulation, suggesting that they can inhibit Se-5HTR. This was supported by their competitive inhibition with the ligand, 5-HT. Bacterial secondary metabolites may be synthesized and released in the bacterium-nematode complex in order to defend immune attack from target insect to compete with other microbes occurring in the insect cadaver and facilitate host nematode development or bacterial quorum sensing [47]. It has been reported phenylethylamides and tryptamides identified from *Xenorhabdus* can act as quorum quenching activators by competitive binding to N-acylated homoserine lactone (AHL) receptor because AHL accumulation drives gene expression of bioluminescence, virulence factor, and biofilm formation in bacteria [60,61]. Xenortides are linear peptides consisting of 2-8 amino acids synthesized from both *Xenorhabdus* and *Photorhabdus* [62]. They are synthesized in insect hosts during infection with putative role in inhibiting prophenoloxidase activation to suppress insect immunity [63]. Xenocycloins produced by *X. bovienii* are cytotoxic to hemocytes of *Galleria melonella* [64]. Nematophin is synthesized by condensation of *α*-keto acid and tryptamine in *X. nematophila.* It possesses a specific antibacterial activity against *Staphylococcus aureus* [65]. GameXPeptides are cyclic pentapeptides widely synthesized in both *Xenorhabdus* and *Photorhabdus.* However, their biological functions remain unclear [66]. This current study showed that these six compound classes could inhibit cellular immune responses by competitive inhibition with 5-HT against Se-5HTR.

A novel phenylethylamide compound, PhX, was found to be highly inhibitory against Se-5HTR. Derivatives of phenylethylamide compound (HB 4) exhibited different inhibitory activities against cellular immune responses mediated by Se-5HTR. Especially, PhX containing methoxy and hexahydropyrrolo[1,2-a]pyrazine-1,4-dione exhibited the highest inhibitory activity. 5-HT receptors have been used for screening for potent insecticides with growth-inhibiting or larvicidal activities against *Pseudaletia separata* [67]. Thus, PhX can be a candidate for this application unless it shows mammalian or non-target species toxicity [68].

## Materials and methods

### Insect rearing and microbial culture

A laboratory strain of *S. exigua* was originated from Welsh onion field (Andong, Korea) and maintained for ~20 years. Larvae of this laboratory strain were reared on an artificial diet [69] at temperature of 25 ± 1°C and relative humidity of 60 ± 10% with a photoperiod of 16:8 h (L:D). Under these conditions, larvae had five instars (‘L1-L5’). Adults were reared with 10% sucrose solution. For immune challenge, *Escherichia coli* Top10 (Invitrogen, Carlsbad, CA, USA) was cultured in Luria-Bertani (LB) medium (BD Korea, Seoul, Korea) in a shaking incubator (200 rpm) at 37°C overnight (16 h).

### Chemicals

Serotonin hydrochloride was purchased from Sigma-Aldrich Korea (Seoul, Korea). It was dissolved in distilled water. SB-269970 (a specific inhibitor to 5-HT receptor subtype 7, 5-HT_7_) was purchased from Cayman Chemical Company (Korea). It was dissolved in desired concentrations with dimethyl sulfoxide (DMSO). Fluorescein isothiocyanate (FITC) [2-(6-hydroxy-3-oxo-3h-xanthen-9-yl)-5-isothiocyanatobenzoic acid] was purchased from Sigma-Aldrich Korea. It was dissolved in DMSO to make a solution at 10 mg/mL. Anticoagulant buffer (ACB) was prepared using 98 mM NaOH, 186 mM NaCl, 17 mM Na_2_EDTA, and 41 mM citric acid at pH 4.5. Phosphate-buffered saline (PBS) was prepared at pH 7.4 with 50 mM sodium phosphate and 0.7% NaCl. Tris-buffered saline (TBS) was prepared using 150 mM NaCl, 50 mM Tris-HCl at pH 7.6. Hank’s balanced salt solution (HBSS) was prepared with the following compositions: 8 g NaCl, 400 mg KCl, 40 mg Na_2_HPO_4_, 60 mg KH_2_PO_4_, 1 g glucose, 140 mg CaCl_2_, 120 mg MgSO_4_, and 350 mg NaHCO_3_ in 1,000 mL distilled H_2_O.

### Bioinformatics to search for 5-HT receptor and sequence analysis

*S. exigua* 5-HT receptor (Se-5HTR) sequence was obtained from GenBank by manual annotation. Briefly, a dopamine receptor sequence (AKR18180.1) of *Chilo suppressalis* was used to screen a transcriptome (SRR1050532) of *S. exigua.* A blast contig (GARL01017386.1) was analyzed for open reading frame (ORF) and the predicted amino acid sequence was used for analysis using BlastP program against GenBank (www.ncbi.nlm.nih.gov). After confirming its high homologies (E value < 10^-20^) with other known insect 5HTRs, the resulting ORF sequence (= Se-5HTR) was deposited at NCBI-GenBank (accession number: MH025798). Sequence alignment was established using Clustal Omega tool (https://www.ebi.ac.uk/Tools/msa/clustalo/) provided by European Bioinformatics Institution. Phylogenetic tree was generated with Neighbor-joining method using Mega6 and ClustalW programs. Bootstrapping values were obtained with 1,000 repetitions to support branch and clustering. Protein domain was predicted using InterPro tool (https://www.ebi.ac.uk/interpro/), pfam (http://pfam.xfam.org), and Prosite (http://prosite.expasy.org/).

### RNA extraction and cDNA preparation

Using Trizol reagent (Invitrogen), total RNAs were extracted from all developmental stages as well as larval tissues (hemocyte, midgut, fat body, and brain) of *S. exigua* according to the instruction of the manufacturer. Numbers of individuals used for RNA extraction for each individual developmental stage were as follows: ~500 eggs, ~20 larvae for L1-L2, ~10 larvae for L3, ~5 larvae for L4, one larva for L5, one pupa, and one adult. L5 larvae were used for RNA extraction from different tissue samples. After extraction, total RNAs were resuspended in nuclease-free water and cDNAs were synthesized from ~1 μg of RNAs using Maxime RT Premix (Intron Biotechnology, Seoul, Korea) according to the manufacturer’s instruction.

### Expression pattern of Se-5HTR by RT-qPCR

A fragment of *Se-5HTR* was amplified with gene-specific primers (5′-CTT TAC CTT CGT GTC TTC TC-3′ and 5′-GGT GTC AGT CTT CTC ATT AC −3′). PCR was performed with 35 cycles of denaturation (94°C, 1 min), annealing (49°C, 1 min), and extension (72°C, 1 min). PCR products were subjected to agarose gel electrophoresis to visually confirm their amplifications. With the same gene-specific primers used in RT-PCR, RT-qPCR was performed in a qPCR machine (CFX Connect™ Real-Time PCR Detection System, Bio-Rad, Hercules, CA, USA) using SYBR Green Realtime PCR Master Mix (Toyobo, Osaka, Japan) under a guideline of Bustin et al. [70]. The amplification used 40 cycles of 15 s at 95°C, 30 s at 60°C, and 45 s at 72°C. After PCR reactions, melting curves from 60 to 95°C were obtained to confirm unique PCR products. A ribosomal protein, RL32, gene was used as a control with primers of 5′-ATG CCC AAC ATT GGT TAC GG-3′ and 5′-TTC GTT CTC CTG GCT GCG GA-3′. Each treatment was independently triplicated. Relative quantitative analysis method (2^-ΔΔCT^) was used to estimate mRNA expression levels of *Se-5HTR*.

### RNA interference (RNAi)

Gene fragment of *Se-5HTR* was amplified from template DNA using gene-specific primers (5’-CTT TAC CTT CGT GTC TTC TC-3’ and 5’-GGT GTC AGT CTT CTC AT-3’) possessing T7 RNA polymerase promoter sequence (5’-TAA TAC GAC TCA CTA TAG GGA GA-3’) at 5’ ends. PCR was performed with 5 cycles of denaturation (94°C, 1 min), annealing (49°C, 1 min), and extension (72°C, 1 min) followed by 30 cycles of denaturation (94°C, 1 min), annealing (60°C, 1 min), and extension (72°C, 1 min) to synthesize DNA template for dsRNA synthesis. Double-stranded RNA (dsRNA) against *Se-5HTR* (‘dsSe-5HTR’) was synthesized using Megascript RNAi kit (Ambion, Austin, TX, USA) following the manufacturer’s instruction. The resulting dsRNA was blended with Metafectene PRO (Biontex, Plannegg, Germany) at 1:1 (v:v) ratio and incubated at 25°C for 30 min for liposome formation. Two microliters of the prepared mixture containing ~900 ng of dsRNA was injected twice into *S. exigua* larval hemocoel using a microsyringe (Hamilton, Reno, NV, USA) equipped with a 26-gauge needle. The first injection was at late L4 stage. It was repeated 12 h afterwards. RNAi efficacy at 0, 24, and 48 h postinjection (PI) in reducing *Se-5HTR* expression was determined by RT-qPCR. At 24 h PI, treated larvae were used for immune challenge experiments. Each treatment was replicated thrice using 10 larvae for each replication.

### Total hemocyte count (THC)

Hemolymph was collected by cutting larval proleg and mixed with ACB (1:10, v/v). Hemocytes were counted using a Neubauer hemocytometer (Superior Marienfeld, Lauda-Königshofen, Germany) under a phase contrast microscope (BX41, Olympus, Tokyo, Japan) at 100× magnification. Heat-killed (90°C, 30 min) *Escherichia coli* (5 × 10^5^ cells/larva) and a test chemical (5-HT or SB-269970) were co-injected into hemocoel through abdominal proleg of L5 larvae in a volume of 5 μL using a 10 μL micro-syringe (Hamilton) after surface-sterilization with 70% ethanol. After 4 h of incubation at 25 ± 2°C, hemolymph of the insect was collected and assessed for THC.

### Hemocyte-spreading analysis

After hemolymph (~150 μL) was collected from five L5 larvae by cutting prolegs, it was mixed with three times volume of ice-cold ACB and incubated on ice for 30 min. ACB-treated hemolymph was then centrifuged at 800 × *g* for 5 min at 4°C. The resulting pellet was resuspended in 500 μL of filter-sterilized TC-100 insect cell culture medium (Welgene, Daegu, Korea). On a glass coverslip placed in a moist chamber, 10 μL of hemocyte suspension was applied and placed in a dark condition. Hemocytes were then fixed with 4% paraformaldehyde (filter-sterilized) at 25°C for 10 min and then washed thrice with filter-sterilized PBS. Cells were then permeabilized with 0.2% Triton-X dissolved in PBS at 25°C for 2 min and washed with PBS. After that, hemocytes were blocked using 10% bovine serum albumin (BSA) dissolved in PBS at 25°C for 10 min and washed again with PBS. Cells were then incubated with FITC-tagged phalloidin in PBS for 60 min and washed thrice with PBS. Hemocytes nuclei were then stained by incubating with 4’,6-diamidino-2-phenylindole (DAPI, 1 μg/mL) (Thermo Fisher Scientific, Rockford, IL, USA) dissolved in PBS and washed thrice with PBS. Hemocytes were then observed under a fluorescence microscope (DM2500, Leica, Wetzlar, Germany) at 400× magnification. Hemocyte-spreading was dictated by appendages of F-actin outward the hemocyte cell boundary.

To assess effect of *Se-5HTR* RNAi on hemocyte-spreading, hemocytes were collected at 24 h after injecting dsSe-5HTR (~900 ng/larva). In separate experiments, 5-HT and 5-HTR inhibitor SB-269970 were co-injected into dsRNA-treated larvae at ratios of 1:10 and 10:1 to check their influence on hemocyte-spreading. At 6 h after co-injection, hemolymph was collected from the treated larva and hemocyte-spreading was observed using the above-mentioned method.

### Nodulation assay

Overnight culture of *E. coli* Top10 bacterial cells was washed with PBS and centrifuged at 1,120 × *g* for 10 min. L5 larvae were used to assess hemocyte nodule formation by immune-challenge with bacterial injection (~1.8 × 10^5^ cells/larva) through the abdominal proleg into larval hemocoel using a microsyringe as previously described. To assess the effect of *Se-5HTR* RNAi on nodule formation, bacterial challenge was performed at 24 h after injecting dsSe-5HTR (~900 ng/larva). To check the influence of chemicals on nodule formation, 2 μg of ketanserin and/or 10 μg of 5-HT was co-injected along with the bacterial suspension. In separate experiment, 1 μg of the bacterial secondary metabolite was co-injected with the bacterial suspension to assess immune suppression activity of the test compound. After an incubation period of 8 h at 25°C, treated insects were dissected under a stereo microscope (SZX9, Olympus, Japan) to count melanized nodule numbers. Each treatment was triplicated independently using five insects for each replication.

### Phagocytosis assay

Preparation of FITC-labeled bacterial cells followed the method described by Harlow and Lane (1998). Briefly, *E. coli* Top10 cells were cultured in 50 mL of Luria-Bertani broth (37°C, 16 h). These bacterial cells were then harvested by centrifuging 1 mL of the cultured broth at 1,120 × *g* for 10 min at 4°C. Bacterial cells (10^5^ cells/mL) were washed twice with TBS and re-suspended in 1 mL of 0.1 M sodium bicarbonate buffer (pH 9.0). In the suspension, 1 μL of 10 mg/mL FITC solution was added and immediately mixed. The mixture was then incubated under darkness with end-over-end rotation at 25°C for 30 min. After the incubation, FITC-tagged bacteria were harvested by centrifugation at 22,000 × *g* for 20 min at 4°C. These bacterial cells were washed thrice with HBSS to remove unbound dye and re-suspended in TBS.

To assess *in vivo* phagocytosis activity, 5 μL of FITC-tagged *E. coli* suspension was injected into the hemocoel of L5 larva through the proleg. To assess the effect of *Se-5HTR* RNAi on phagocytosis activity, bacteria were injected at 24 h after injecting dsSe-5HTR (~900 ng/larva). To check the influence of a specific 5-HT_7_ receptor inhibitor on phagocytosis, 2 μg of SB-269970 was co-injected with the tagged bacterial suspension. To determine the effect of secondary metabolite on phagocytosis, 1 μg of bacterial secondary metabolite was co-injected with tagged bacterial suspension. After 15 min of incubation, treated larvae were surface-sterilized using 70% ethanol. With a pair of scissors, the proleg was cut to collect hemolymph sample (~50 μL) in 150 μL of cold ACB with gentle shaking of the tube to mix hemolymph and ACB thoroughly. Hemocyte monolayers were made using 50 μL of hemocyte suspension (~5 × 10^3^ cells) and left in a moist chamber for 15 min for hemocytes to settle and attach to the glass surface. After the incubation period, monolayers were washed with TBS to remove plasma. These monolayers were overlaid with 1% trypan blue dye solution to quench non-phagocytosed bacterial cells. After 10 min, monolayers were washed again with TBS and then fixed with 1.5% glutaraldehyde solution to observe under a fluorescence microscope at 400× magnification. Hemocytes undergoing phagocytosis were counted from a total of 100 hemocytes observed from different areas of each slide. Each observation was triplicated with three different slides.

### Preparation of organic extracts from bacterial culture broth

*X. nematophila* K1 (Xn) and *P. temperate temperata* ANU101 (Ptt) bacteria were cultured in TSB at 28°C for 48 h. Culture broths were centrifuged at 12,500 × *g* for 30 min and supernatants were used for subsequent fractionation. To obtain ethyl acetate extract, the same volume (1 L) of ethyl acetate was mixed with the supernatant and separated into organic and aqueous fractions. Ethyl acetate extract (‘EAX’) was dried using a rotary evaporator (Sunil Eyela, Seongnam, Korea) at 40°C. The resulting extract (0.2 mg) was obtained from 1 L cultured broth and resuspended with 5 mL of methanol. The aqueous phase was then combined with 1 L of butanol. Butanol extract (‘BX’) was also dried using the rotary evaporator at 40°C and the resulting extract (0.2 mg) was resuspended with 5 mL of methanol.

### Secondary bacterial metabolites – biological activity against immune responses

Secondary metabolites (37 samples, S4 Fig) derived from *Xenorhabdus* and *Photorhabdus* cultures were from the Bode lab compound collection named ‘HB’ compounds. Their biological activities for suppressing *S. exigua* immunity were then determined. Individual chemicals were dissolved in DMSO, diluted into desired concentrations with DMSO, and stored at −20°C.

For hemocyte nodulation inhibition assay, overnight culture of *E. coli* bacterial cells was washed with PBS. Test compound (1 μg/larva) was injected into larval hemocoel along with the bacterial suspension (~1.8 × 10^5^ cfu/larva) using a microsyringe as previously described. Insects were then incubated at 25°C for 8 h. After the incubation period, insects were dissected and nodule numbers were counted as described above. Each treatment was triplicated independently using five insects for each replication. For phagocytosis inhibition assay, 5 μL of FITC-tagged *E. coli* suspension along with 1 μg of test compound was injected into L5 larval hemocoel. After 15 min of incubation period, hemocytes undergoing phagocytosis were counted as previously described.

### Secondary bacterial metabolites – competitive assay with 5-HT

To determine effects of bacterial secondary metabolites on nodulation and phagocytosis, 10 potent chemicals were selected based on their common inhibitory activity on both phagocytosis and nodulation and their median inhibition concentration (IC_50_) values were calculated. Percentages of phagocytosis against increasing concentrations of HB chemicals were calculated and their IC_50_ values were calculated using Probit analysis (https://probitanalysis.wordpress.com). To assess a competitive inhibitory activity between test compound and 5-HT, HB chemicals were injected in different doses (0, 0.01, 0.1, 1 and 10 μg/larva) along with a fixed 5-HT concentration (1 μg/larva) and FITC-tagged bacteria (500 cells/larva). In a separate experiment, different doses of 5-HT were injected into the larvae with a fixed HB compound content (1 μg/larva). At 15 min after bacterial injection, hemocytes from treated larvae were collected in ACB and phagocytosis assay was performed as described above.

### Chemical derivatives and their inhibitory activities against Se-5HTR

Based on potent phenylethylamide (PEA) HB compounds, 45 additional PEA samples (S5 Fig) were obtained from Korea Chemical Bank of the Korea Research Institute of Chemical Technology (KRICT). Derivative compounds were dissolved in DMSO, diluted into desired concentrations with DMSO, and stored at −20°C. These PEA chemicals were then tested for their abilities to suppress nodule formation and phagocytosis in *S. exigua* as described above. For nodulation inhibition assay, each chemical (150 ng/larva) was injected into larval hemocoel along with bacterial suspension (~1.8 × 10^5^ cfu/larva). Each treatment was triplicated independently using five insects for each replication. For phagocytosis inhibition assay, 5 μL of FITC-tagged *E. coli* suspension along with 150 ng of each chemical was injected into L5 larval hemocoel of *S. exigua*. After incubating for 15 min, hemocytes undergoing phagocytosis were counted using previously described method.

### Chemical synthesis of PhX

A potent chemical was designed as PhX ((S)-2-(1,4-dioxohexahydropyrrolo[1,2-*a*]pyrazin-2(*1H*)-yl)-*N*-(4-methoxyphenethyl)acetamide) and chemically synthesized according to a method described in S6 Fig.

### Statistical analysis

All studies were triplicated independently. Results are expressed as mean ± standard error. Results were plotted using Sigma plot (Systat Software, San Jose, CA, USA). Means were compared by least squared difference (LSD) test of one-way analysis of variance (ANOVA) using POC GLM of SAS program [71] and discriminated at Type I error = 0.05.

### Funding

This work was supported by a grant (No. 2017R1A2133009815) of the National Research Foundation (NRF) funded by the Ministry of Science, ICT and Future Planning, Republic of Korea. This work was also supported by a project (No SI1808, Development of crop protection agents leading the future market). The chemical library used in this research was provided by Korea Chemical Bank (http://www.chembank.org) of the Korea Research Institute of Chemical Technology (KRICT).

## Acknowledgments

We appreciate Youngim Song for ordering and arranging materials and Malsook Cho for rearing insects. Work in the Bode lab was supported by the LOEWE Center Translational Biodiversity Genomics funded by the state of Hesse, Germany.

## Author Contributions

**Conceptualization**: Yonggyun Kim.

**Formal analysis**: Ariful Hasan, Yonggyun Kim.

**Investigation**: Hyun-Suk Yeom, Jaewook Ryu, Helge B. Bode, Yonggyun Kim.

**Project administration**: Yonggyun Kim.

**Supervision**: Hyun-Suk Yeom, Jaewook Ryu, Helge B. Bode, Yonggyun Kim.

**Writing – original draft**: Ariful Hasan, Yonggyun Kim.

**Writing – review & editing**: Ariful Hasan, Hyun-Suk Yeom, Jaewook Ryu, Helge B. Bode, Yonggyun Kim.

## Supplementary data

**S1 Fig.**
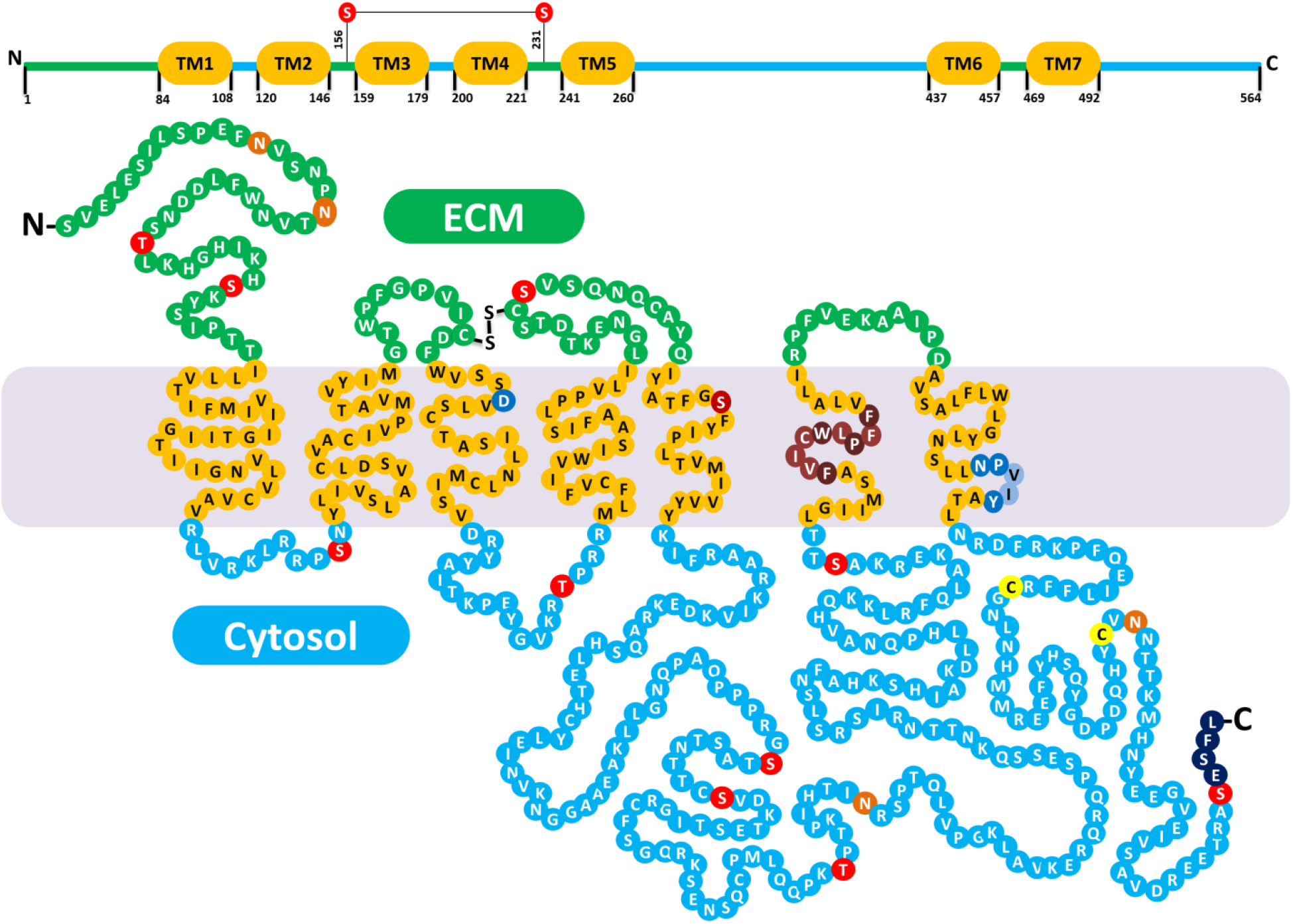
Putative domain and motif structures of Se-5HTR. Putative seven transmembrane domains (TM1-TM7) are primarily marked by dark yellow bars and circles. Extracellular and cytosolic regions are denoted by green and light blue regions, respectively. Potential N-glycosylation sites and phosphorylation sites are marked by orange and red circles, respectively. Aspartic acid residue (dark blue circle) in TM3 and serine residue (light brown circle) in TM5 are putative residues that might chemically interact with 5-HT. The unique consensus sequence motif (PXXXWXPXF, dark brown circles) in aminergic receptors is conserved in TM6. The motif (NPXXY, dark blue circles) is conserved in TM7 like other GPCRs. Two possible post-translational palmitoylation cysteine residues (light yellow circles) and a PDZ-domain binding motif (ESFL, black circles) are also present in the C-terminal. Conserved motifs were determined using InterPro tool (https://www.ebi.ac.uk/interpro/) and Prosite (http://prosite.expasy.org/) whereas other residues and motifs were predicted using several tools from DTU bioinformatics.

**S2 Fig.**
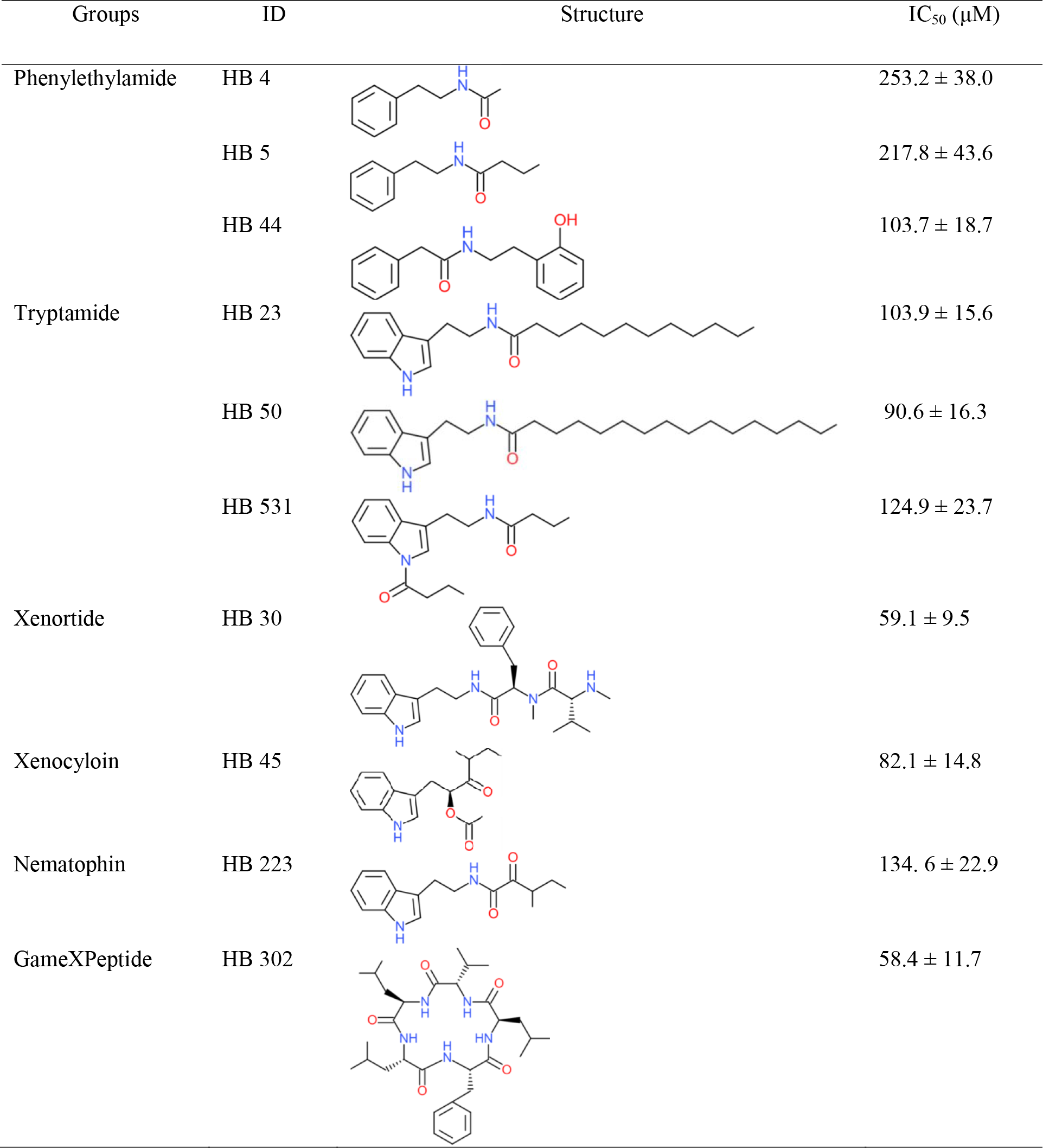
Potent screened chemicals from HB compounds with their chemical structures and respective median inhibitory concentrations (IC_50_). HB chemicals were injected in different doses (0, 0.01, 0.1, 1, and 10 μg/larva) along with a fixed 5-HT concentration (1 μg/larva) and FITC-tagged bacteria (500 cells/larva). After 15 min of treatment, hemocytes from treated larvae were collected in ACB followed by phagocytosis assay as described above. Percentages of phagocytosis against HB chemical treatment with increasing concentrations were calculated. Their IC_50_ values were determined using Probit analysis (https://probitanalysis.wordpress.com).

**S3 Fig.**
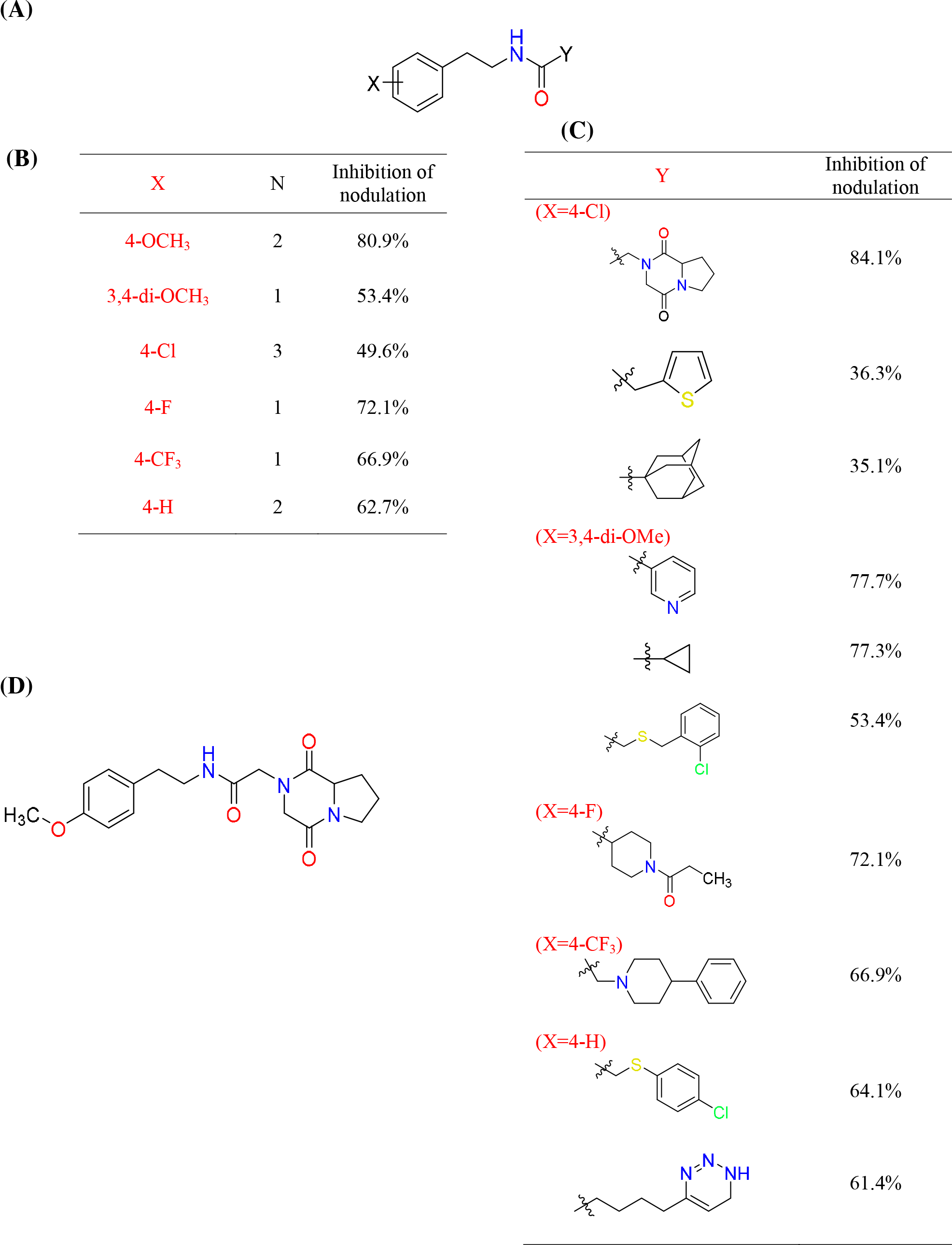
Designing a potent chemical inhibitor from phenylehtylamide (PEA) derivatives. Derivatives of PEA were tested for their nodulation inhibition percentages and sorted by their X and Y groups. Most potent residues inhibiting nodulation were selected and a hypothetical most potent PEA chemical was designed. (A) Core PEA structure with two variable hypothetical residues (X and Y). X belongs to the residue group attached with para position of the phenyl ring whereas Y belongs to the residue group linked to the amide group. (B) Comparative analysis between X residue groups with their mean percent inhibition of nodulation. (C) Comparative analysis between Y residue groups with their percent inhibition of nodulation. (D) A hypothetical PEA chemical compound structure having the most potent inhibition capability.

**S4 Fig.**
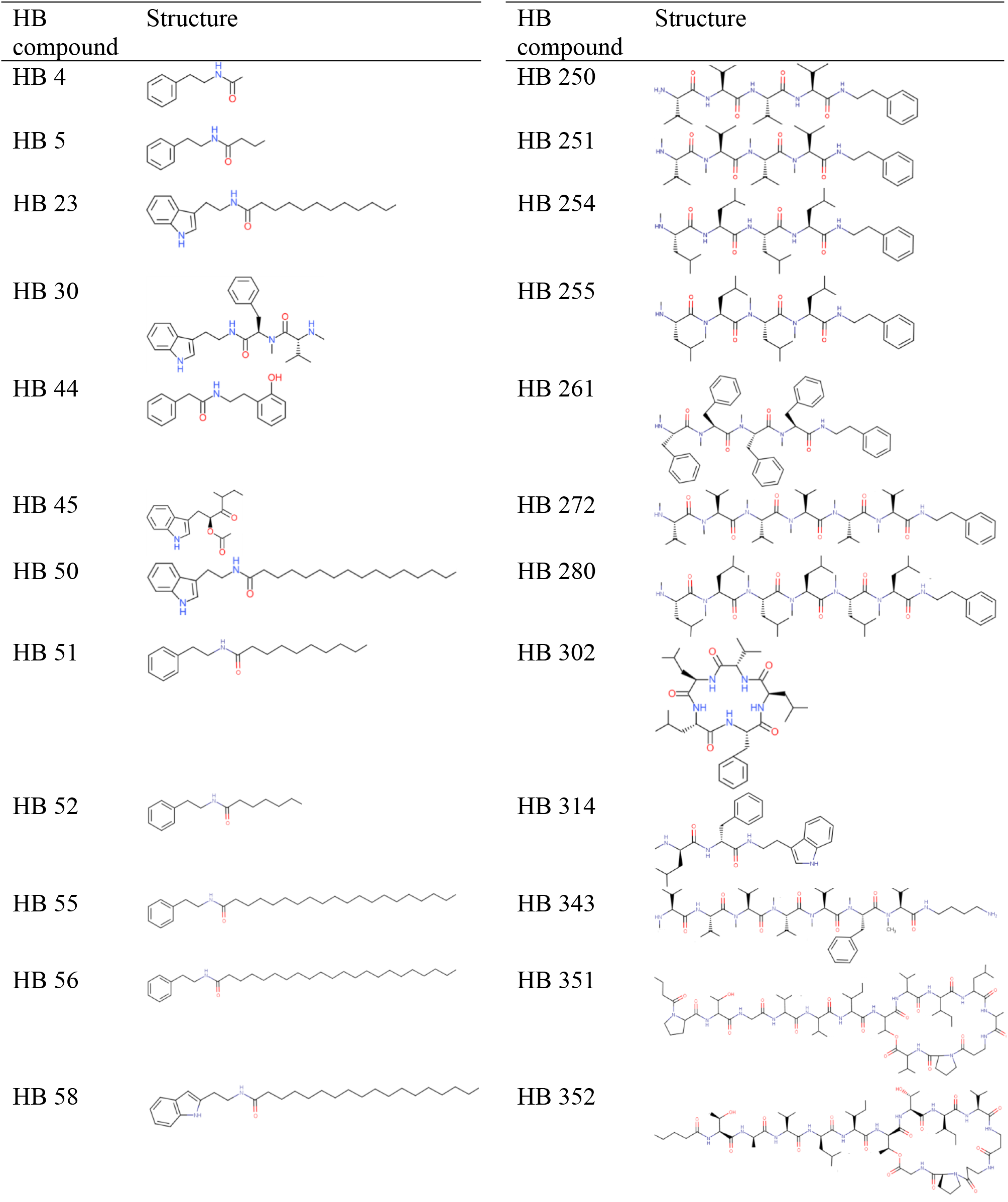
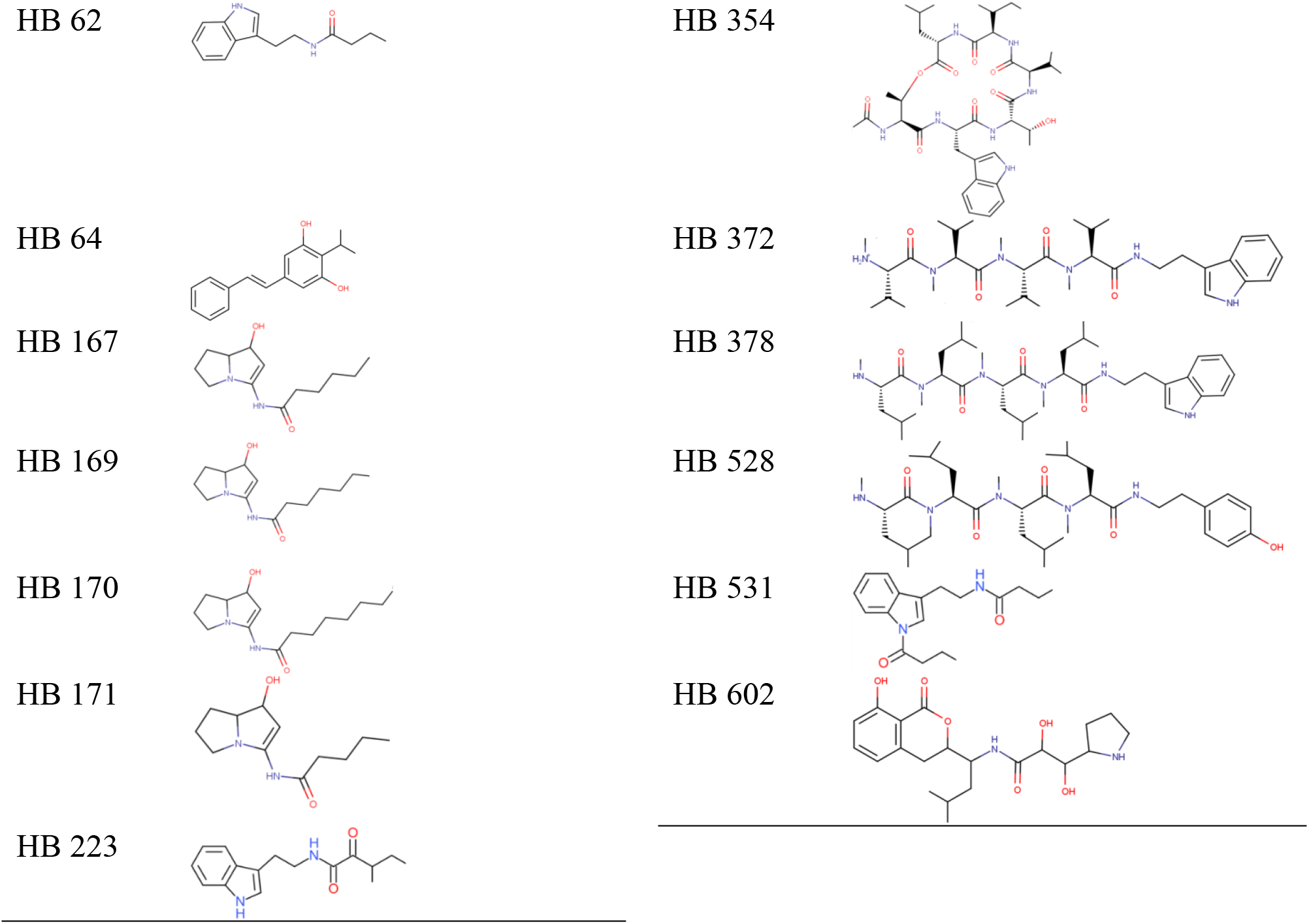
Bacterial secondary metabolites (37 chemicals) derived from *Xenorhabdus* and *Photorhabdus*.

**S5 Fig.**
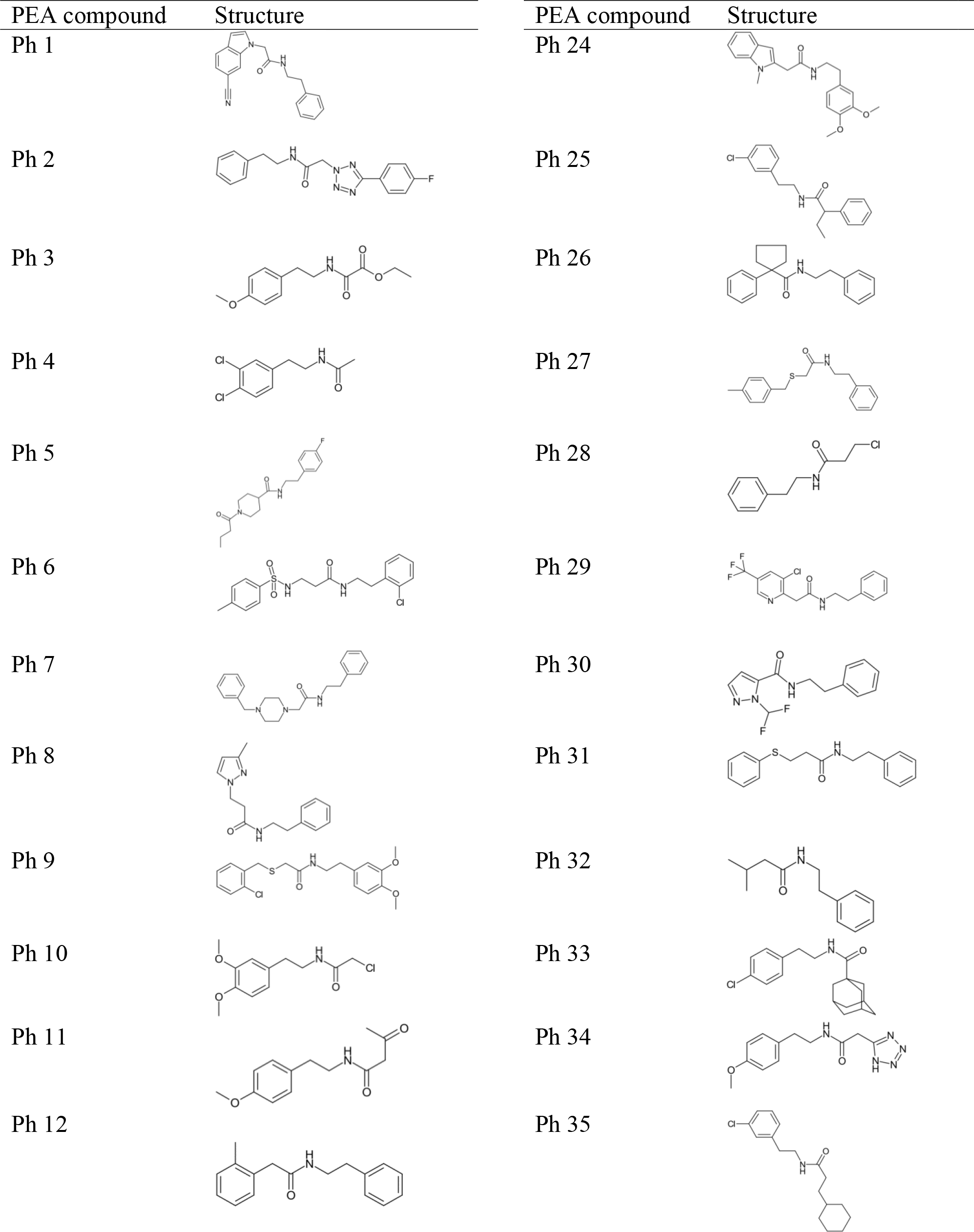
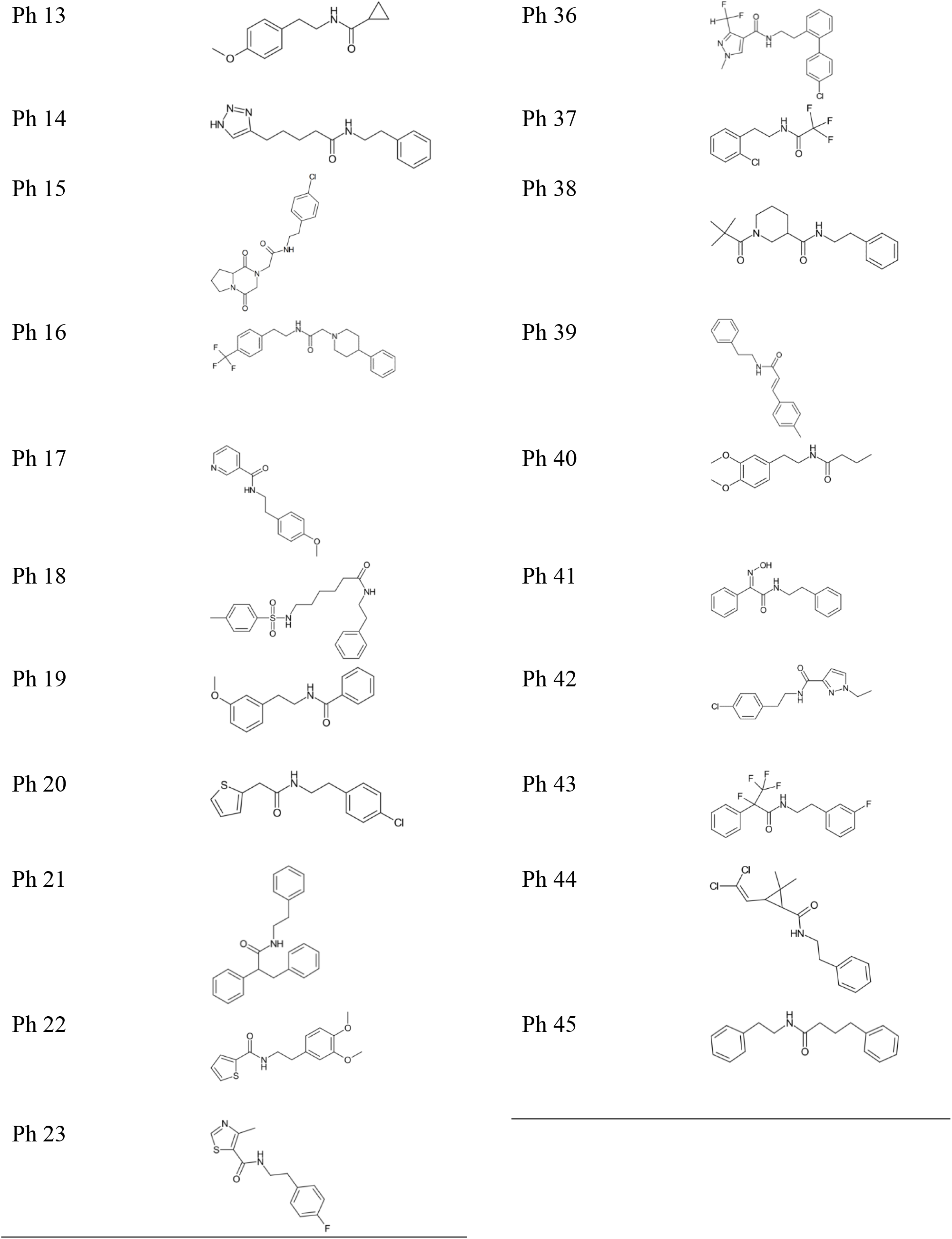
Phenylethylamide (PEA) derivatives (45 chemicals) based on HB 44, a bacterial metabolite

**S6 Fig.**
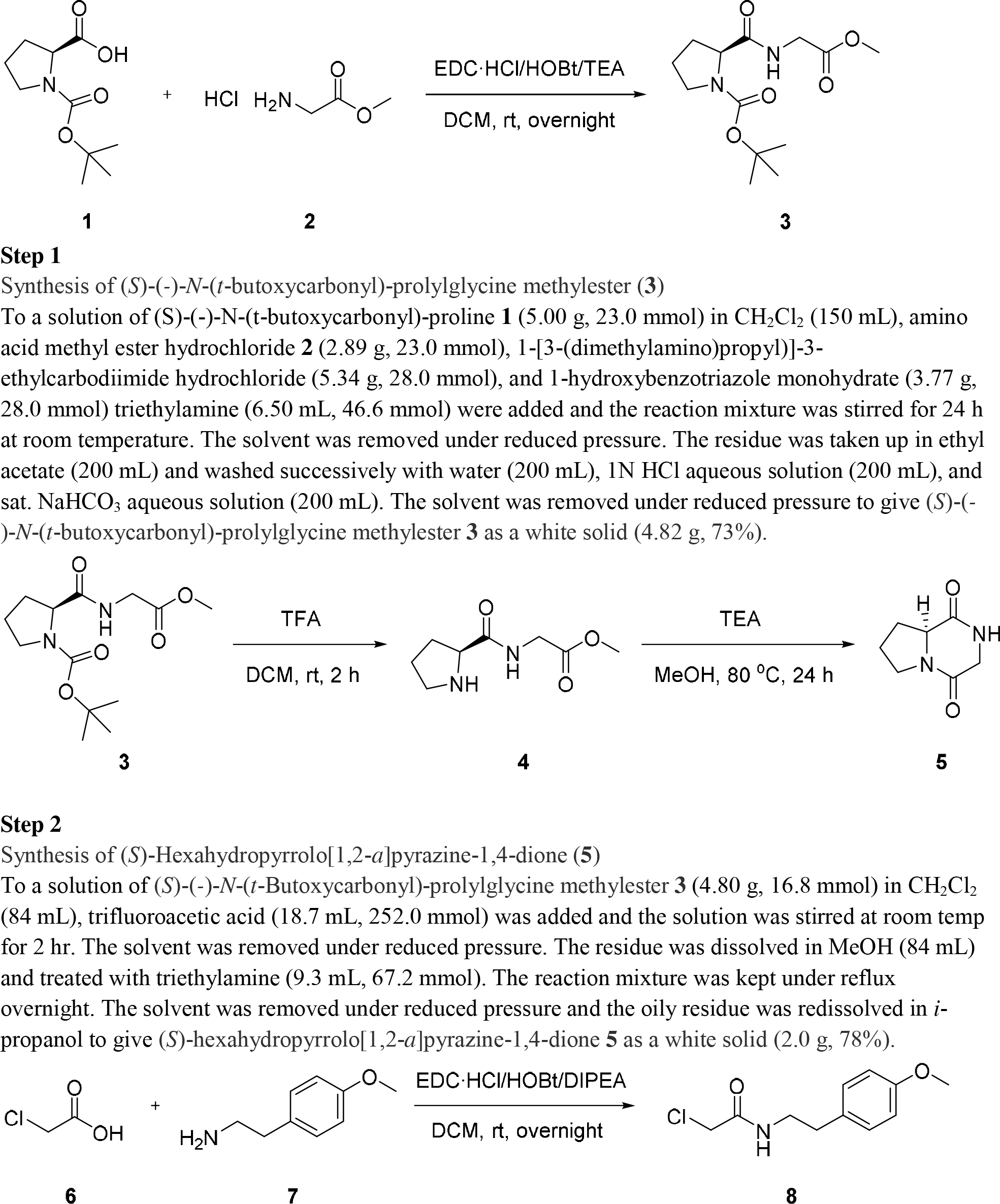
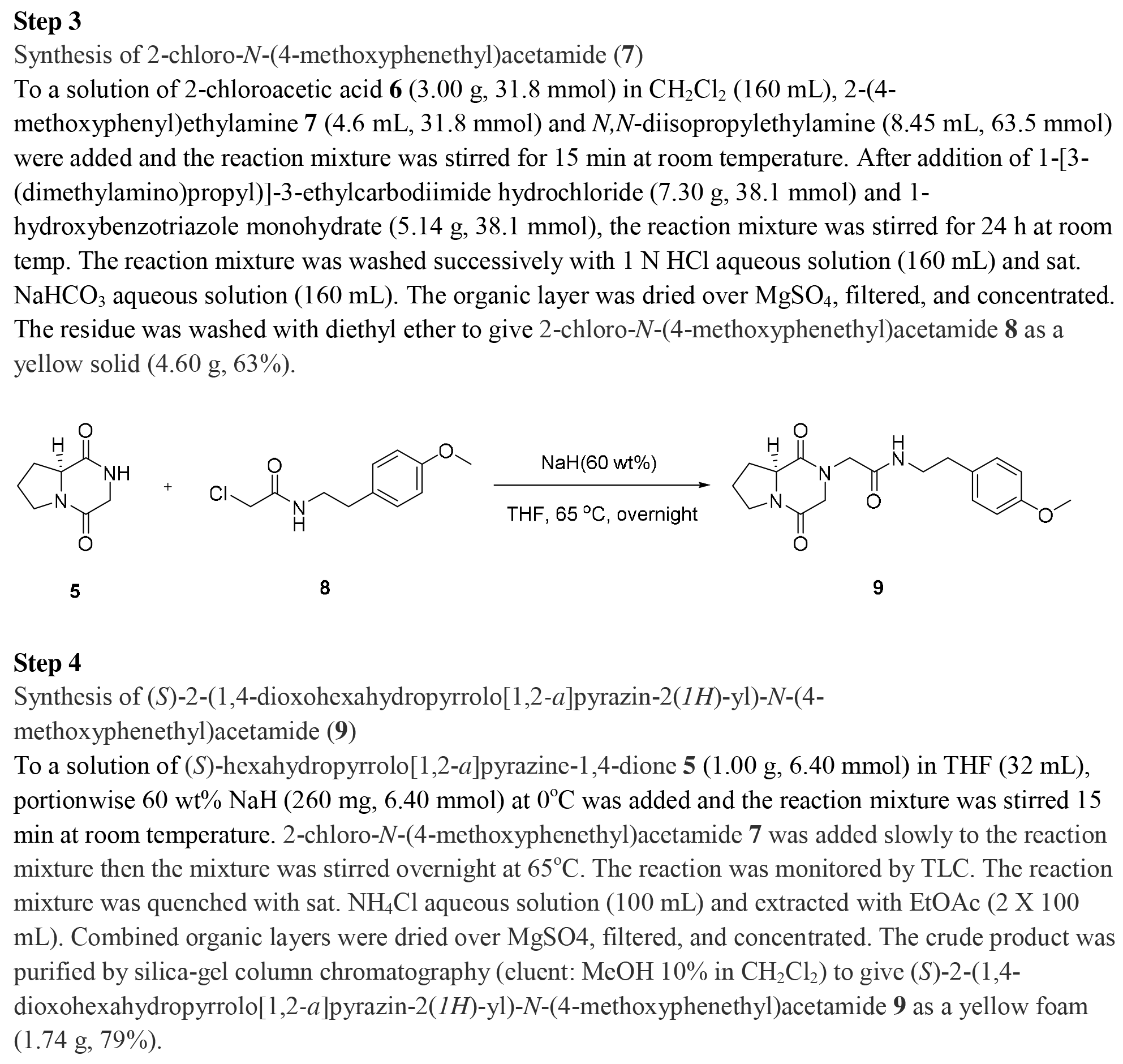
Chemical synthesis of PhX ((*S*)-2-(1,4-dioxohexahydropyrrolo[1,2-*a*]pyrazin-2(*1H*)-yl)-*N*-(4-methoxyphenethyl)acetamide)

## References

1. Roshchina VV. 2010. New trends and perspectives in the evolution of neurotransmitters in microbial, plant, and animal cells. Adv Exp Med Biol. 2016;874:25–77.

2. Lovenberg W, Weissbach H, Udenfriend S. 1962. Aromatic L-amino acid decarboxylase. J Biol Chem. 1962;237:89–93.

3. Hufton SE, Jennings IG, Cotton RG. Structure and function of the aromatic amino acid hydroxylases. Biochem. J. 1995;311:353–66.

4. Roberts KM, Fitzpatrick PF. 2013. Mechanisms of tryptophan and tyrosine hydroxylase. IUBMB Life 2013;65:350–357.

5. Zhang X, Beaulieu JM, Sotnikova TD, Gainetdinov RR, Caron MG. Tryptophan hydroxylase-2 controls brain serotonin synthesis. Science. 2004;305:217.

6. Gershon MD, Tack J. The serotonin signaling system: from basic understanding to drug development for functional GI disorders. Gastroenterology. 2007;132:397–414.

7. Richtand NM, McNamara RK. Serotonin and dopamine interactions in psychosis prevention. Prog Brain Res. 2008;172:141–53.

8. Monti JM. Serotonin control of sleep-wake behavior. Sleep Med Rev. 2011;15:269–81.

9. Švob Štrac D, Pivac N, Mück-Šeler D. The serotonergic system and cognitive function. Transl Neurosci. 2016;7:35–49.

10. Seuwen K, Pouysségur J. Serotonin as a growth factor. Biochem. Pharmacol. 1990;39:985–90.

11. Li N, Wallén NH, Ladjevardi M, Hjemdahl P. Effects of serotonin on platelet activation in whole blood. Blood Coagul Fibrinolysis 1997;8:517–23.

12. Hayashi K, Fujita Y, Ashizawa T, Suzuki F, Nagamura Y, Hayano-Saito Y. Serotonin attenuates biotic stress and leads to lesion browning caused by a hypersensitive response to *Magnaporthe oryzae* penetration in rice. Plant J. 2016;85:46–56.

13. Majeed ZR, Abdeljaber E, Soveland R, Cornwell K, Bankemper A, Koch F, et al. Modulatory action by the serotonergic system: behavior and neurophysiology in *Drosophila melanogaster*. Neural Plast. 2016;2016:1–23.

14. Huser A, Eschment M, Güllü N, Kan C, Böpple K, Pankevych L. et al. Anatomy and behavioral function of serotonin receptors in *Drosophila melanogaster* larvae. PLoS One 2017;12:e0181865.

15. Becnel J, Johnson O, Luo J, Nässel DR, Nichols CD. The serotonin 5-HT_7Dro_ receptor is expressed in the brain of *Drosophila*, and is essential for normal courtship and mating. PLoS One 2011;6:e20800.

16. Yuan Q, Lin F, Zheng X, Sehgal A. Serotonin modulates circadian entrainment in *Drosophila*. Neuron. 2005;47:115–27.

17. Nichols CD. 5-HT_2_ receptors in *Drosophila* are expressed in the brain and modulate aspects of circadian behaviors. Dev Neurobiol. 2007;67:752–63.

18. Yuan Q, Joiner WJ, Sehgal A. A sleep-promoting role for the *Drosophila* serotonin receptor 1A. Curr. Biol. 2006;16:1051–62.

19. Neckameyer WS, Coleman CM, Eadie S, Goodwin SF. Compartmentalization of neuronal and peripheral serotonin synthesis in *Drosophila melanogaster*. Genes Brain Behav. 2007;6:756–69.

20. Dierick HA, Greenspan RJ. Serotonin and neuropeptide F have opposite modulatory effects on fly aggression. Nat. Genet. 2007;39:678–82.

21. Kaplan DD, Zimmermann G, Suyama K, Meyer T, Scott MP. A nucleostemin family GTPase, NS3, acts in serotonergic neurons to regulate insulin signaling and control body size. Genes Dev. 2008;22:1877–93.

22. Qi YX, Huang J, Li MQ, Wu YS, Xia RY, Ye GY. Serotonin modulates insect hemocyte phagocytosis via two different serotonin receptors. Elife 2016;5:e12241.

23. Python F, Stocker RF. Immunoreactivity against choline acetyltransferase, gamma-aminobutyric acid, histamine, octopamine, and serotonin in the larval chemosensory system of *Drosophila melanogaster*. J. Comp. Neurol. 2002;453:157–67.

24. Dasari S, Cooper RL. Direct influence of serotonin on the larval heart of *Drosophila melanogaster*. J. Comp. Physiol. B 2006;176:349–57.

25. Rodriguez MVG, Campos AR. Role of serotonergic neurons in the *Drosophila* larval response to light. BMC Neurosci. 2009;10:66.

26. Barnes, N.M., Sharp, T., 1999. A review of central 5-HT receptors and their function. Neuropharmacology 38, 1083–1152.

27. Nichols DE, Nichols CD. Serotonin receptors. Chem Rev. 2008;108:1614–41.

28. Roth BL. The serotonin receptors: from molecular pharmacology to human therapeutics. Humana Press, Totowa, New Jersey. 2006.

29. Thompson AJ, Lummis SCR. 5-HT_3_ Receptors. Curr. Pharm. Des. 2006;12:3615–30.

30. Watanabe T, Sadamoto H, Aonuma H. Identification and expression analysis of the genes involved in serotonin biosynthesis and transduction in the field cricket *Gryllus bimaculatus*. Insect Mol Biol. 2011;20:619–35.

31. Ono H, Yoshikawa H. Identification of amine receptors from a swallowtail butterfly, *Papilio xuthus* L.: cloning and mRNA localization in foreleg chemosensory organ for recognition of host plants. Insect Biochem Mol Biol. 2004;34:1247–56.

32. Vleugels R, Lenaerts C, Baumann A, Vanden BJ, Verlinden H. Pharmacological characterization of a 5-HT_1_-type serotonin receptor in the red flour beetle, *Tribolium castaneum*. PLoS One 2013;8:e65052.

33. Troppmann B, Balfanz S, Baumann A, Blenau W. Inverse agonist and neutral antagonist actions of synthetic compounds at an insect 5-HT_1_ receptor. Br. J. Pharmacol. 2010;159:1450–62.

34. Guo X, Ma Z, Kang L. Serotonin enhances solitariness in phase transition of the migratory locust. Front. Behav Neurosci. 2013;7:129.

35. Qi YX, Xia RY, Wu YS, Stanley D, Huang J, Ye GY. Larvae of the small white butterfly, *Pieris rapae*, express a novel serotonin receptor. J Neurochem. 2014;131:767–77.

36. Qi YX, Jin M, Ni XY, Ye GY, Lee Y, Huang J. Characterization of three serotonin receptors from the small white butterfly, *Pieris rapae*. Insect Biochem. Mol. Biol. 2017;87:107–16.

37. Thamm M, Balfanz S, Scheiner R, Baumann A, Blenau W. Characterization of the 5-HT_1A_ receptor of the honeybee (*Apis mellifera*) and involvement of serotonin in phototactic behavior. Cell Mol. Life Sci. 2010;67:2467–79.

38. Schlenstedt J, Balfanz S, Baumann A, Blenau W. Am5-HT_7_: molecular and pharmacological characterization of the first serotonin receptor of the honeybee (*Apis mellifera*). J Neurochem. 2006;98:1985–98.

39. Berridge MJ, Patel N. Insect salivary glands – stimulation of fluid secretion by 5-hydroxytryptamine and adenosine-3’, 5’-monophosphate. Science. 1968;162:462–3.

40. Berridge MJ. The role of 5-hydroxytryptamine and cyclic AMP in the control of fluid secretion by isolated salivary glands. J Exp Biol. 1970;53:171–86.

41. Vanhoenacker P, Haegeman G, Leysen JE. 5-HT_7_ receptors: current knowledge and future prospects. Trends Pharmacol. Sci. 2000;21:70–7.

42. Molaei G, Lange AB. The association of serotonin with the alimentary canal of the African migratory locust, *Locusta migratoria*: distribution, physiology and pharmacological profile. J Insect Physiol. 2003;49:1073–182.

43. Baines D, DeSantis T, Downer RGH. Octopamine and 5-hydroxytryptamine enhance the phagocytic and nodule formation activities of cockroach (*Periplaneta americana*) haemocytes. J Insect Physiol. 1992;38:905–14.

44. Kim K, Madanagapol N, Lee D, Ki Y. Octopamine and 5-hydroxytryptamine mediate hemocytic phagocytosis and nodule formation via eicosanoids in the beet armyworm, *Spodoptera exigua*. Arch. Insect Biochem. Physiol. 2009;90:162–76.

45. Kim GS, Kim Y. Up-regulation of circulating hemocyte population in response to bacterial challenge is mediated by octopamine and 5-hydroxytryptamine via Rac1 signal in *Spodoptera exigua*. J. Insect Physiol. 2010;56:559–66.

46. Kim Y, Ji D, Cho S, Park Y. Two groups of entomopathogenic bacteria, *Photorhabdus* and *Xenorhabdus*, share an inhibitory action against phospholipase A2 to induce host immunodepression. J Invertebr Pathol. 2005;89: 258–64.

47. Shi YM, Bode HB. Chemical language and warfare of bacterial natural products in bacteria-nematode-insect interactions. Nat Prod Rep 2018;35:309–35.

48. Tobias NJ, Shi YM, Bode HB. Refining the natural product repertoire in entomopathogenic bacteria. Trends Microbiol. 2018;26:833–40.

49. Barak LS, Tiberi M, Freedman NJ, Kwatra MM, Lefkowitz RJ, Caron MG. A highly conserved tyrosine residue in G protein-coupled receptors is required for agonist-mediated ß2-adrenergic receptor sequestration. J Biol Chem 1994;269:2790–5.

50. Ji TH, Grossmann M, Ji I. G protein-coupled receptors. I. Diversity of receptor-ligand interactions. J Biol Chem. 1998;273:17299–302.

51. Huang ES. Construction of a sequence motif characteristic of aminergic G protein-coupled receptors. Protein Sci. 2003;12:1360–67.

52. Park J, Stanley D, Kim Y. Rac1 mediates cytokine-stimulated hemocyte spreading via prostaglandin biosynthesis in the beet armyworm, *Spodoptera exigua*. J Insect Physiol. 2013;59:682–9.

53. Venkatakrishnan AJ, Deupi X, Lebon G, Tate CG, Schertler GF, Babu MM. Molecular signatures of G-protein-coupled receptors. Nature 2013;494:185–94.

54. Ho BY, Karschin A, Branchek T, Davidson N, Lester HA. The role of conserved aspartate and serine residues in ligand binding and in function of the 5-HT_1A_ receptor: A site-directed mutation study. FEBS Lett. 1992;312:259–62.

55. Shapiro DA, Kristiansen K, Weiner DM, Kroeze WK, Roth BL. Evidence for a model of agonist-induced activation of 5-hydroxytryptamine 2A serotonin receptors that involves the disruption of a strong ionic interaction between helices 3 and 6. J. Biol. Chem. 2002;277:11441–9.

56. Qanbar R, Bouvier M. Role of palmitoylation/depalmitoylation reactions in G-protein-coupled receptor function. Pharmacol. Ther. 2003;97:1–33.

57. Romero G, von Zastrow M, Friedman PA. Role of PDZ proteins in regulating trafficking, signaling, and function of GPCRs: means, motif, and opportunity. Adv Pharmacol. 2011;62:279–314.

58. Guseva D, Wirth A, Ponimaskin E. Cellular mechanisms of the 5-HT_7_ receptor-mediated signaling. Front. Behav. Neurosci. 2014;8:306.

59. Choi C, Helfman DM. The Ras-ERK pathway modulates cytoskeleton organization, cell motility and lung metastasis signature genes in MDA-MB-231 LM2. Oncogene 2014;33:3668–76.

60. Papenfort K, Bassler BL. Quorum sensing signal-response systems in Gram-negative bacteria. Nat. Rev. Microbiol. 2016;14:576–88.

61. Bode E, He Y, Vo TD, Schultz R, Kaiser M, Bode HB. Biosynthesis and function of simple amides in *Xenorhabdus doucetiae*. Environ. Microbiol. 2017;19:4564–75.

62. Reimer D, Nollmann FI, Schultz K, Kaiser M, Bode HB. Xenortide biosynthesis by entomopathogenic *Xenorhabdus nematophila*. J Nat Prod. 2014;77:1976–80.

63. Bloudoff K, Schmeing TM. Structural and functional aspects of the nonribosomal peptide synthetase condensation domain superfamily: discovery, dissection and diversity. Biochim Biophys Acta 2017;1865:1587–604.

64. Proschak A, Zhou Q, Schöner T, Thanwisai A, Kresovic D, Dowling A, et al. Biosynthesis of the insecticidal xenocyloins in *Xenorhabdus bovienii*. Chembiochem. 2014;15:369–72.

65. Li J, Chen G, Webster JM. Nematophin, a novel antimicrobial substance produced by *Xenorhabdus nematophilus* (Enterobactereaceae). Can J Microbiol. 1997;43:770–3.

66. Nollmann FI, Dauth C, Mulley G, Kegler C, Kaiser M, Waterfield NR. et al. Insect-specific production of new GameXPeptides in *Photorhabdus luminescens* TTO1, widespread natural products in entomopathogenic bacteria. Chembiochem. 2015;16:205–8.

67. Cai M, Li Z, Fan F, Huang Q, Shao X, Song G. Design and synthesis of novel insecticides based on the serotonergic ligand 1-[(4-Aminophenyl) ethyl]-4-[3-(trifluoromethyl) phenyl] piperazine (PAPP). J Agric Food Chem. 2009;58:2624–9.

68. Verlinden H, Vleugels R, Broeck JV. Serotonin, serotonin receptors and their actions in insects. Neurotransmitter. 2015;2:e314

69. Shrestha S, Stanley D, Kim Y. PGE2 induces oenocytoid cell lysis via a G protein-coupled receptor in the beet armyworm, *Spodoptera exigua*. J. Insect Physiol. 2011; 57:1568–76.

70. Bustin SA, Benes V, Garson JA, Hellemans J, Huggett J, Kubista M, et al. The MIQE guidelines: minimum information for publication of quantitative real-time PCR experiments. Clin. Chem. 2009;55:4.

71. SAS Institute. SAS/STAT user’s guide. SAS Institute, Inc., Cary, NC. 1989.

